# Functional Differentiation of Type II and Type I Collagen Articular Models in Synovial Fluid Film Formation and Recombinant Lubricin Retention

**DOI:** 10.64898/2026.05.22.726594

**Authors:** Diego R. Jaramillo Pinto, Nicolas L. Mendoza, Syeda Tajin Ahmed, Yidan Wen, Lenka Vitkova, Spencer M. Witt, Kaleb A. Cutter, Ummay Honey, Matthew J. Paszek, Heidi L. Reesink, Lawrence J. Bonassar, Kevin De France, Roberto C. Andresen Eguiluz

## Abstract

Collagen type II (Col-II) and collagen type I (Col-I) are major components of articular cartilage present at different ratios at its surface. Understanding how each of these components mediates the assembly of molecular films derived from synovial fluid (SF), the native lubricant of synovial joints, is critical to explain the loss of mechanical performance in pathological conditions, guide the design of biomaterial implants meant to be in contact with SF, and develop molecular therapies to restore SF properties. This work demonstrates that Col-II articular surface model assists in scaffolding of full SF-derived films, while Col-I model lacks SF film scaffolding capabilities. However, when Col-II and Col-I are exposed to recombinant lubricin (rLub) alone, the major boundary lubricant in SF, both adsorbed and retained similar amounts. These insights, deduced from quartz crystal microbalance with dissipation, diffuse reflectance circular dichroism, and atomic force microscopy, reveal possible mechanisms underlying the loss of mechanical performance of synovial joints in pathology, where Col-I becomes the major collagenous component of the articular cartilage surface, as well as considerations for designing functional biomaterial implants. Furthermore, this work reinforces the idea of rLub as an intra-articular osteoarthritis therapy with the ability to bind to Col-II and Col-I, irrespectively.

**GRAPHICAL ABSTRACT:** 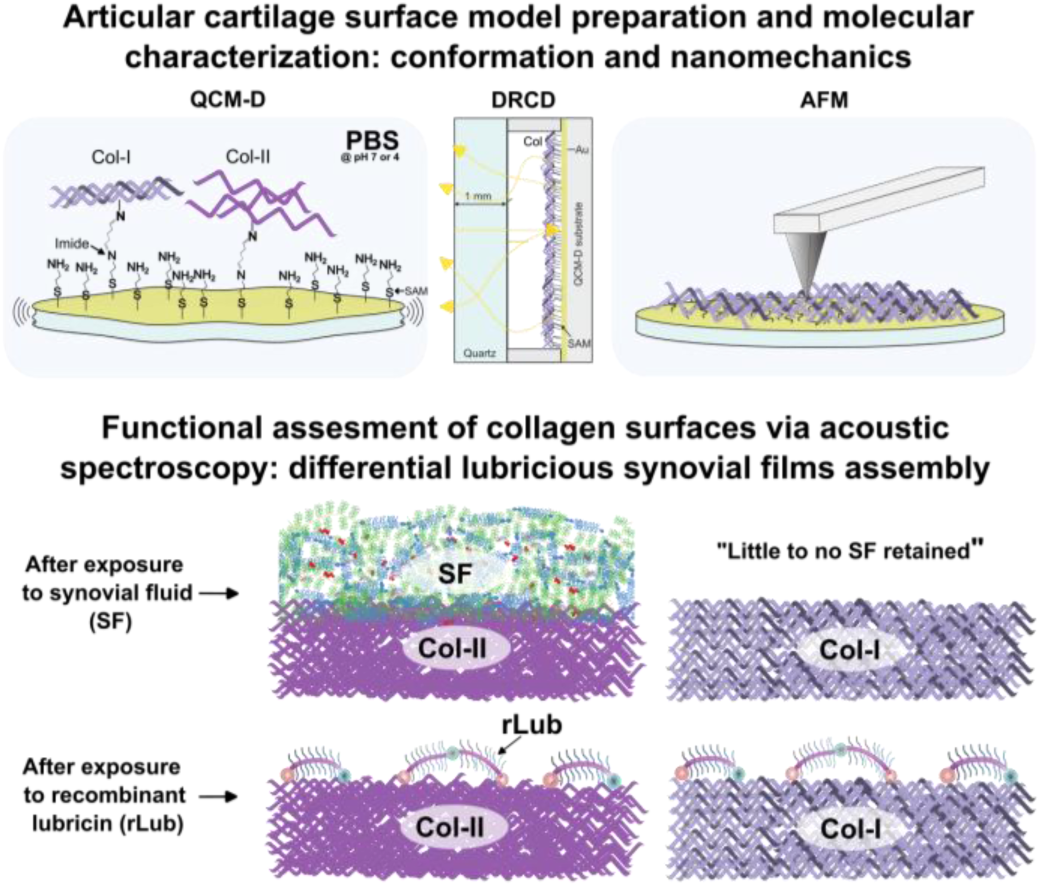

## INTRODUCTION

Synovial joints are defined by their mobility, where bone surfaces are covered by articular cartilage that enables low-friction movement and repetitive mechanical loading, facilitated by synovial fluid (SF) lubrication. At the interface between the surface of the articular cartilage extracellular matrix (AC-ECM) and SF, a thin film of SF components existing as a gel-like layer dissipates loads and reduces wear, damage, and friction forces in boundary lubrication mode^[1,2]^. The degree of hydration at this interface has been proposed to be, in part, responsible for the tribological robustness of synovial joints^[3–5]^, along with synergistic interactions between glycosaminoglycans (GAGs, *e.g.,* hyaluronic acid, HA), phospholipids (PLs, *e.g.*, phosphatidylcholine, PC), and glycoproteins (GP, *e.g.,* lubricin, Lub), whose individual roles (or low-order mixtures) have been widely investigated^[4,6–20]^. Another key mechanism proposed for boundary lubrication involves steric-entropic repulsive forces that result from the flexibility of coiled molecules resisting conformational restriction upon confinement, as has been demonstrated for Lub alone and Lub mixed with HA ^[18,21]^. Thus, recombinant Lub (rLub) has been proposed as a key molecule with potential to treat synovial joint diseases, such as osteoarthritis ^[22–25]^.

While these studies have shaped our understanding of synovial joint nanomechanics, fully capturing SF synergies on a molecular level is essential to enable the efficient design of the next generation of intra-articular osteoarthritis therapies. Collagen type II (Col-II) is the main component of healthy AC-ECM (Figure 1 (A)), while collagen type I (Col-I) is the major collagen component in damaged cartilage or fibrocartilaginous repair tissue (Figure 1 (B))^[26,27]^. Some studies have shown that SF components, such as Lub, can bind to Col-II at the molecular level, forming anti-adhesive and wear-protecting layers ^[28,29]^. Also, solid phase binding assays have shown that rLub can bind to Col-II and fibronectin (FN), another molecular component of the AC-ECM^[30]^.

**Figure 1.**
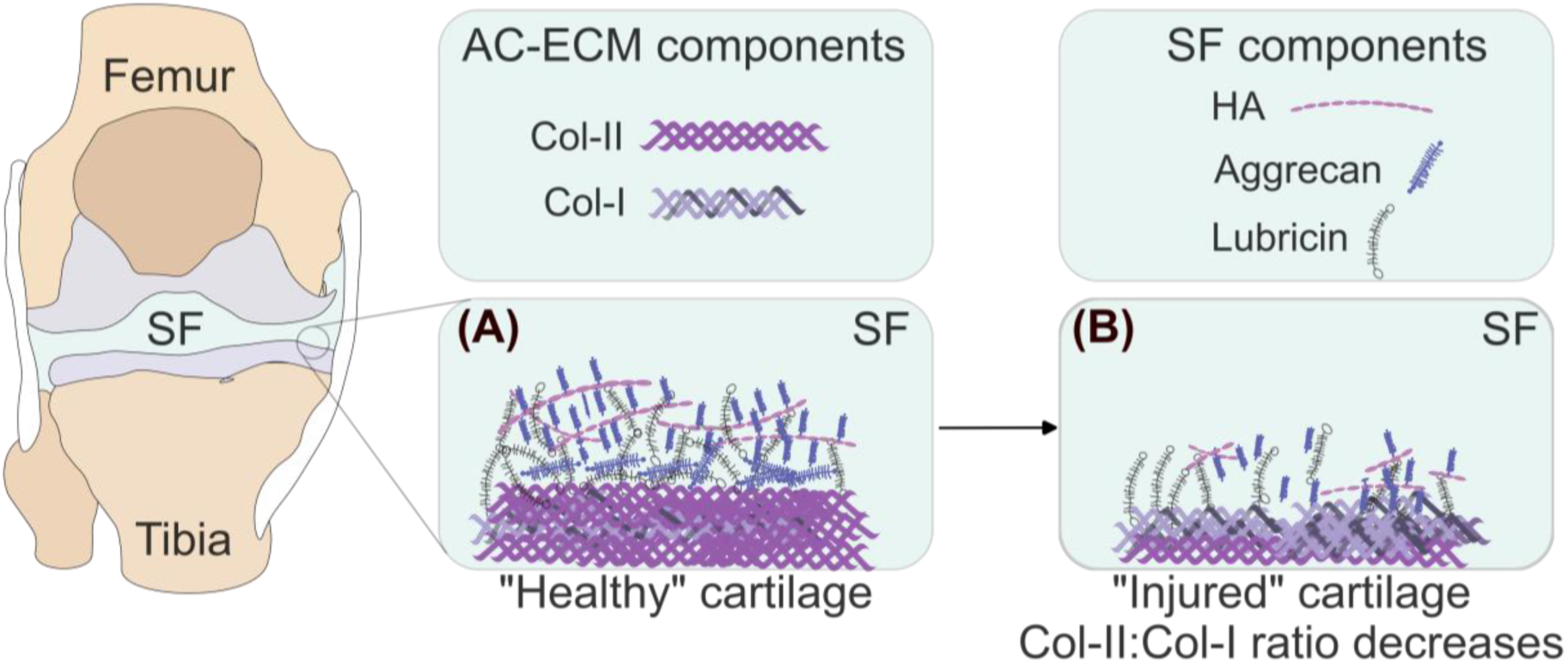
Schematic of a knee (prototypical synovial joint) and the regions in which this study focuses, *i.e.*, the interface between SF and AC-ECM. (A) and (B) represent “healthy” and “injured” cartilages respectively, considering only the collagenous components of the AC-ECM and some SF components.

Efforts have been made to fabricate matrices of Col-II^[31]^, Col-I^[32]^, and mixtures of both^[33]^ in order to support cartilage repair^[34]^, screening of preclinical drugs^[35]^, and understanding how both of these collagens interact with key SF components at the molecular level ^[28,29,36]^. However, the roles of Col-II and Col-I in the supramolecular assembly of a film derived from the totality of SF components, as well as the resulting nanomechanical properties of that film, such as normal (load-carrying) and lateral (friction) compliance, remain unexplored. Furthermore, it is well established that the constituents of extracellular matrices appear in a range of molecular conformation distributions, which are affected and altered by disease and mechanical stimuli^[37–44]^. How AC-ECM alterations, such as molecular conformations of collagens (*e.g.* II or I), and surface morphologies, may affect the adsorption of SF components, and thus SF film assembly and mechanics, is yet another open question. This is important to guide effective pathophysiological treatments, including the design and application of biomaterials for cartilage repair and synthetic SFs, based on, for example, rLub.

Col-II and Col-I are fibril-forming collagens and consist of three α-polypeptide chains that form a right-handed triple helix, generating a collagen molecule^[45]^. While mostly similar, they differ in key molecular features that give each its own characteristics. On one hand, both collagen molecules have a molecular weight of ∼ 280-300 kDa, a contour length of ∼ 300 nm, and a diameter of ∼ 1.5 nm^[45,46]^. However, Col-II is a homotrimer consisting of three α_1_(II) polypeptide chains ^[45]^, whereas Col-I is a heterotrimer consisting of two α1(I) and one α2(I) polypeptide chains, encoded by different genes. α_1_(I), α_1_(II) and α_2_(I) have different amino acid sequences, more specifically, different interruptions of the typical Gly-X-Y amino acid sequence^[47]^. These chains assemble onto triple helices to create collagen molecules, which then assemble with a staggered arrangement into fibrils whose length and diameter *in vitro* are in the order of microns and nanometers, respectively^[48–50]^. In the AC-ECM, one of the principal roles of collagen molecules is to provide mechanical integrity.

In solution, the stability of collagen triple helices, and thus their tertiary structure is strongly dependent on the environmental conditions, such as pH ^[49,51,52]^. In this work, molecular films of Col-II and Col-I were formed by controlling their bulk conformations via solution pH adjustment, to either pH ∼ 7 or pH ∼ 4, followed by chemical immobilization onto NH_2_-terminated self-assembled monolayers (SAMs) on gold-coated quartz crystals via glutaraldehyde activation^[28,53]^. This process yielded immobilized collagen films with different nano-morphologies and nano-mechanics as evidenced by measurements of hydrated film mass density, Young’s modulus, film elastic compliance, film morphology, and film molecular conformation. To characterize these films, quartz crystal microbalance with dissipation (QCM-D), atomic force microscopy (AFM) in liquid environment, and diffuse reflectance circular dichroism (DRCD) were used. Furthermore, the efficacy of these films to retain SF and rLub was assessed via QCM-D.

Results revealed differences in the assembly of SF-derived films on Col-II compared to Col-I surfaces. Col-II retained significantly larger amounts of SF components than Col-I. On the other hand, all Col-II and Col-I films retained similar amounts of rLub, suggesting that the binding affinity of rLub is similar for Col-II and Col-I under the studied conditions. These results give insight into how altered AC-ECM loses SF supramolecular assembly when Col-II is replaced by Col-I, with possible implications in poor load-bearing capabilities^[13]^ and consequently loss of mechanical performance. On the other hand, our findings suggest that the tested rLub could be explored for therapeutic use in injured cartilage, or for supplementation of lubrication-deficient SF.

## MATERIALS AND METHODS

### Materials

Phosphate buffer saline (PBS) (Gibco), ethanol (ACROS, absolute, 200 Proof, ≥99.5%), sodium dodecyl sulfate (SDS) (MP Biomedicals, ultrapure, ≥99%) were used as received. Base-piranha solution was prepared by mixing ammonium hydroxide (Chemsavers, 28-30% pure), and hydrogen peroxide (PERDROGEN^TM^ by Honeywell, MDL # MFCD00011333, ≥30% w/w stabilized) and water as follows: 6:1:1 by volume of water: ammonium hydroxide: hydrogen peroxide. Gold-coated silica quartz crystals were purchased from Quartz PRO with a resonance frequency of 5 MHz and a diameter of 14 mm. For gold surface functionalization, cysteamine hydrochloride (Sigma, ≥98% titration) and glutaraldehyde (Sigma, 25% aqueous solution, Grade I) were used. Ultrapure water was collected from a Thermo-Fisher UV water purification system, with a resistivity of 18.2 MΩ·cm.

### Col-II and Col-I

Bovine collagen type II (Col-II, Sigma-Aldrich, Southern Biotech), bovine collagen type I (Col-I, Gibco) were used for collagen film formation. Col-II and Col-I solutions were stored as-purchased at 4 °C. The working concentration of 50 µg/mL was prepared with PBS with a pH of 7.4 (referred to as PBS 7 or PBS at pH 7), or 4 (referred to as PBS 4 or PBS at pH 4). PBS with a pH of 4 was prepared by adding tiny acetic acid (0.1 M) droplets and was used within a month.

### Bovine synovial fluid

Bovine SF was stored at −80 °C until use. To prepare working aliquots, frozen SF was thawed and centrifuged for 5 min at 6000 rpm to remove cell debris and large tissue aggregates, followed by a 25% v/v dilution in PBS as a thinning step to prevent QCM-D lines from clogging. Aliquots were stored at −80 °C until the day of experiment. This is referred to as dilute SF, or dSF in the text.

### Recombinant lubricin

Recombinantly-produced equine lubricin (rLub) was manufactured using codon scrambling^[54]^, resulting in a maximum stock concentration of 20 mg/ml. The stock was diluted to 5 mg/ml in PBS and friction tested using a custom pin-on-plate tribometer^[55]^ to confirm lubricity (details are provided in the SI, with results shown in Figure S1). For QCM-D testing, solutions were stored at –20 °C and further diluted to a working concentration of 100 µg/mL on the day of the experiment.

All experiments using collagen, SF, and rLub were conducted in accordance with the procedures and guidelines of the Institutional Biosafety Committee at the University of California, Merced.

### Functionalization of substrates with self-assembled monolayers for QCM-D

Clean gold-coated crystals with a 5 MHz fundamental resonance frequency were rinsed with 2 wt% SDS, ultrapure water, and ethanol, respectively, and the process was repeated three times. After rinsing, surfaces were dried with a gentle stream of N_2_, followed by an oxygen plasma cleaning step (Harrick Plasma, PDC-32G-115V) for 2 min and stored in Petri dishes for immediate use.

Clean crystals were incubated in 0.088 M ethanolic solution of cysteamine hydrochloride at 4^°^ C overnight. This process creates amine-terminated thiol-self-assembled monolayers on the gold surfaces and helps retain collagen, preventing collagen desorption due to Vroman effects^[56]^ when exposing the collagen films to synovial fluid. The next day, surfaces were gently rinsed with ethanol to remove excess unbound NH_2_-terminated molecules and then dried under N_2_ gas. Then, surfaces were incubated in 25% aqueous glutaraldehyde solution at room temperature for 30 minutes. This allowed imine formation between the aldehyde group of glutaraldehyde and the amine group of cysteamine hydrochloride (immobilization schematic shown in Figure 2(A)). After incubation, substrates were rinsed with water and then dried with an N_2_ stream again. Functionalized surfaces were immediately loaded into the QCM-D fluidic cell.

**Figure 2.**
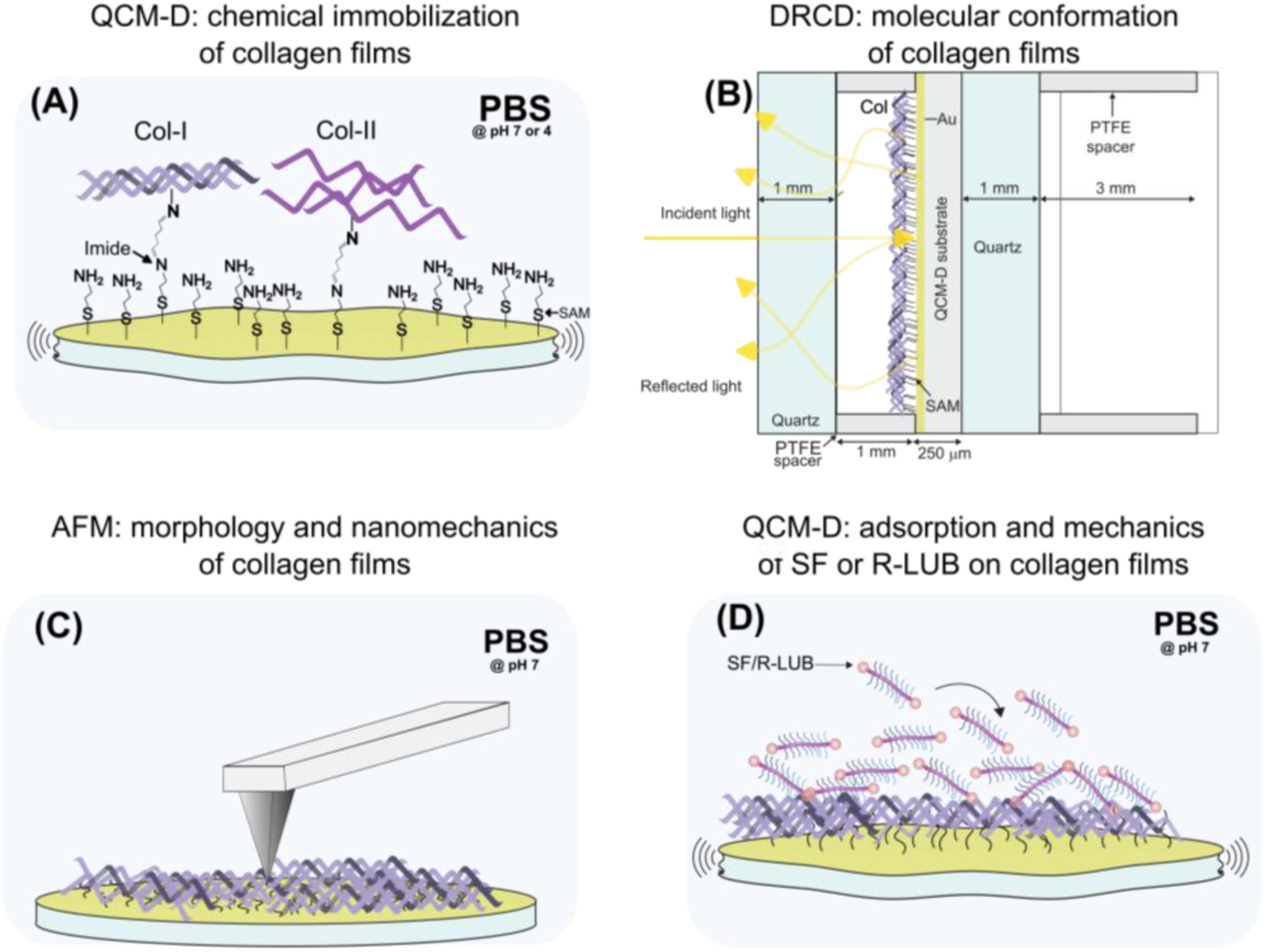
Schematic summarizing experimental approaches and techniques, which include surface functionalization and characterization via QCM-D, DRCD and AFM. (A) Chemical immobilization of Col-II/Col-I to amine-terminated SAM surfaces through glutaraldehyde activation, in PBS at pH 7 or 4. (B) Assessing molecular conformation of collagen films via DRCD. (C) Collagen film morphology and mechanics investigated via AFM in liquid environment measurements of the collagen films. (D) Quantifying adsorption of SF/rLub to the collagen surfaces in PBS at pH 7 via QCM-D.

After each use, crystals were cleaned in base piranha by submersion for ∼30 seconds at 60 °C. Next, crystals were rinsed with 2% SDS, water, and ethanol multiple times, then dried with a gentle stream of dry N_2_, finishing with a short plasma cleaning step. Each crystal was reused on average 5 times without loss of sensitivity.

### Quartz Crystal Microbalance with Dissipation (QCM-D)

QCM-D measurements were performed using a QCM-I mini (Micro Vacuum, Hungary) and a Q-Sense microbalance (Biolin, Sweden), using AT-cut quartz crystals with 5 MHz fundamental frequency coated with 50 nm of gold. All measurements were performed at 25 ^°^C controlled by an incorporated Peltier unit. The outlet channels of the QCM-I mini and Q-Sense were connected to peristaltic pumps (BQ80S Microflow Variable-Speed Peristaltic Pump and ISMATEC, respectively), which maintained a constant flow rate. All the QCM-D data presented in the main text come from the steady state.

The QCM-D experimental sequence began with a PBS (at pH 7 or 4) baseline (*i.e*., until a frequency dispersion of 2 Hz/hr or less over two minutes was achieved). Next, collagen solutions at concentrations ranging from 10 to 100 µg/ml were flowed by applying suction with a peristaltic pump until the fluidic chamber was saturated. After saturation, the peristaltic pump was stopped, and changes in frequency signal and dissipation were collected until a steady state was achieved. This step is named “collagen pre-wash” in the text. PBS 7 or PBS 4 was used to wash loosely bound Col-II and Col-I molecules, and the signal was collected until a steady state was reached. This step is denoted as “Col-II/Col-I post-wash”. For experiments in which the collagen films were deposited at pH 4, a second PBS wash was performed at pH 7. In the text, data from this step is denoted as “pH 4 to 7”. Up to this point, the experimental sequence was the same for both types of experiments, collagen films exposed to dSF and rLub. Next, for the rLub experiments, 100 µg/ml of rLub in PBS was flowed until a steady state was achieved (“rLub pre-wash”), followed by a PBS wash (“rLub post-wash”). On the other hand, for the dSF experiments, a blocking step with 10 mg/ml BSA in PBS at pH 7 was performed (“BSA pre-wash”), followed by a PBS wash at pH 7 (“BSA post-wash”). After this step, dSF was flowed and data was collected until a steady state was achieved (“dSF pre-wash”). The experiments concluded with a wash with PBS at pH 7 (“dSF post-wash”).

The last two minutes of the steady state values (Δ*f* and Δ*D*) of Col-II or Col-I pre- and post-wash, dSF pre- and post-wash, and rLub pre- and post-wash were used to calculate the average mass densities, *m*_f_, and thin film elastic compliances, 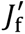, as described below. To calculate *m*_f_ of rLub adsorption on collagen films, frequency changes between the rLub pre- and -post-washes steps were compared against the last PBS wash performed on the respective collagen film. For the case of dSF adsorption, the comparison was between dSF pre- and post-washes and BSA post-wash step. This approach allowed us to determine the areal mass of adsorbed dSF and retained dSF after wash, without accounting for the precursor films on functionalized gold surfaces. For shear film compliance, because deconvoluting the contribution of each layer is not straightforward, each condition was compared with the baseline, i.e., a stacked system, for cases with rLub and dSF components adsorbed to the collagen films.

Each reported condition consists of at least three independent measurements, reported as the average ± standard error of the mean in SI tables, unless otherwise stated.

Control experiments with only Col-II or Col-I adsorption on bare gold (without any functionalization), rLub adsorption experiments on bare gold and on SAM-functionalized gold were also performed. dSF adsorption controls were omitted from this report, as these experiments were previously published by our group elsewhere^[53]^.

### QCM-D data analysis and statistics

QCM-D data were analyzed using a combination of OriginLab Pro and Microsoft Excel. In this report, QCM-D data is shown as box-and-whisker plots, displaying the mean (solid line), median (broken line), lower and upper quartiles (box), and minimum/maximum values excluding outliers with a coefficient of 1.5 (whiskers). Each presented data point was utilized for calculations of the mean and SEM. A data point is an independent measurement corresponding to the average of two minutes of data collected on the steady state of each condition, using pyQCM-BraTaDio[55]. Statistical differences were assessed using a combination of two-way ANOVA, in OriginLab, and unpaired two-sample Student-s *t*-test. For *t*-test, two groups were considered significantly different from each other if the *p*-value was 0.05 or lower (one asterisk between them on the plots). When *p* < 0.01, two asterisks are shown, and when *p* < 0.001, three asterisks are shown. The *t*-tests were performed with a Python[56] script based on Scipy[57] and tkinter, written with the assistance of AI.

### Modeling of QCM-D data

Mass density (*m*_f_) and shear elastic compliance 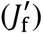 of precursor (collagen) films and of adlayers (dSF/rLub) were calculated with the software pyQCM-BRATADIO^[57]^. *m*_f_, using the Sauerbrey model, can be estimated as:

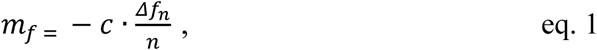

where *c* is the crystal mass sensitivity, typically 17.7 ng·cm^−2^·Hz^−2^ for a 5 MHz crystal, and *Δf*_n_ is the change in frequency for the overtone *n.* A linear regression was used to obtain the slope of the change in frequency versus all the available overtones, typically from the 3^rd^ to the 13^th^, as described in ^[53]^. Shear modulus (*G*_f_) is also reported as the inverse of shear compliance on a second y-axis. The Sauerbrey thickness, *d*_f_, was estimated as follows: an effective dry mass density for the collagen films was calculated from^[58]^:

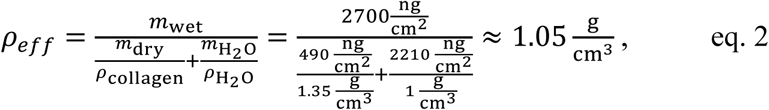

using the value of dry mass density for Col-II in PBS at pH 7 on gold, at 50 µg/ml, reported by Yuan et al.^[29]^ (*m*_dry_), as 490 ng/cm^2^, and of *m*_wet_ of 2700 ng/cm^2^, as obtained via QCM-D and reported for Col-II on bare Au (results in SI). A similar calculation for the case of Col-I was performed, using the Col-I value of 340 ng/cm^2^ on Au at 48 µg/ml reported by Yuan et al.^[36]^ obtaining values even closer to 1 g/cm^3^, *i.e.*, to a film density dominated by water. Thus, the value obtained from eq. 2 was chosen to be used as *ρ*_*eff*_ across all films, assuming that it can be true for hydrated protein films with a 10% error or less. From this, a Sauerbrey thickness *d*_f_ in nanometers can be estimated as:

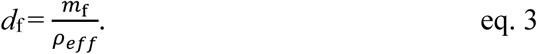

The shear elastic compliance, 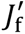 was calculated using:

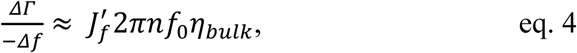

as derived by Du and Johannsmann^[59]^, where *f*_0_ is the fundamental resonance frequency, *η*_*bulk*_is the viscosity of the media, and Δ𝛤 is the change in half-band half-width. It is related to changes in dissipation *D* as follows:

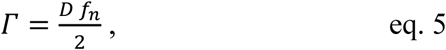

with *f*_n_ =*n*·*f*_0_ as the resonance frequency of the *n*^th^ overtone. In this approximation, it is assumed that the density of the bulk fluid (PBS) and the immobilized/adsorbed films are similar, which, based on Eq. 2, is likely to be true within a 10% error. Representative QCM-D traces for changes in frequency and dissipation of each condition are shown in Figure S2 (dSF experiments) and Figure S3 (rLub experiments).

### Immobilization of collagens for DRCD and AFM

Immobilization of Col-I and Col-II for DRCD and AFM analysis was performed outside the QCM-D fluidic chamber as follows: a clean QCM-D crystal was functionalized with a SAM and activated with glutaraldehyde, as for QCM-D measurements. After the glutaraldehyde activation step, the crystals were rinsed with ultrapure water, followed by complete immersion in a clean glass jar containing 50 µg/mL protein solution at the corresponding pH for 2 hr at 4 °C, followed by a PBS wash at their respective pH, and ending with a PBS 7 wash for the case of pH 4 conditions.

### DRCD

DRCD spectra were collected on a circular dichroism spectrometer J-1500 (JASCO, Japan) operating in diffuse reflectance mode under a nitrogen atmosphere. Samples were placed in a sample holder and immersed in PBS (pH 7). A total of 10 spectra accumulations were collected and averaged, with a digital integration time of 4 s, a bandwidth of 4 nm, and a scanning speed of 100 nm/min. Spectra were baseline-corrected, smoothed with a Savitzky-Golay filter (polynomial order 1, frame length 9)^[60]^, and zero-endpoint corrected at the long-wavelength limit of each window. Data is presented as mean residue ellipticity (*MRE*), with units of mdeg ⋅ m^2^ ⋅ mol^−1^ ⋅ residue^−1^, calculated from:

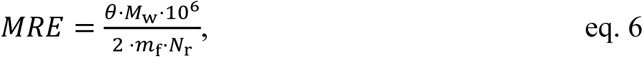

where *θ* is the measured ellipticity in millidegrees, *m*_f_ is the surface-specific mean Sauerbrey mass in ng/cm², *M*_w_ is the protein molecular weight in g/mol and *N*_r_ is the total residue count per molecule. These values are shown in Table S1.

For collagen samples with the spectra exhibiting the canonical triple-helix band structure, the ratio between the magnitudes of the positive peak at 222 nm and the negative peak at 197 nm^[61]^ is reported as:

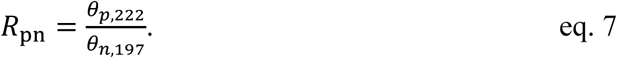

Because the *R*_pn_ is a ratio of two ellipticities at two wavelengths, it is mathematically invariant under any scalar multiplier of the spectrum (effective concentration, path length, film thickness) and is therefore in principle well-defined for surface-confined DRCD films. It should be noted that *R*_pn_ from DRCD is not numerically comparable to published transmission-CD *R*_pn_ values because DRCD and transmission CD use different baseline-subtraction conventions (bare-substrate / zero-endpoint vs. solvent blank), and the additive residual baseline does not cancel in a ratio. *R*_pn_ is therefore reported for relative comparison within the present dataset only. For conditions where the canonical triple-helix was not present, the comparisons were limited to spectral shape and *MRE* magnitudes at the diagnostic wavelengths.

### CD

The bulk collagen samples were measured in PBS solution at a protein concentration of 0.01mg/mL, measured in transmission using quartz cuvettes with the pathlength of 1 mm. The samples were measured as the average of three spectra accumulations from 250-190 nm at 25 °C, and were subsequently baseline and zero-endpoint corrected, and converted to *MRE*. *R*_pn_ was also calculated according to eq. 7.

### AFM

A Nanowizard Pure AFM (Bruker, USA) was used for imaging immobilized protein films immersed in PBS at pH 7. ScanAsyst-Fluid probes (Bruker, USA), with a nominal normal spring constant of 0.7 N/m were used. Cantilevers were V-shaped, gold-coated on the backside. QI mode was used to obtain height maps of collagen films, as it relies on a fixed tip-sample force setpoint, allowing imaging of soft surfaces with minimal damage. For the images of collagen films with rLub, contact mode imaging was performed, to visualize possible differences in the tip-sample interactions when scanning on rLub molecules. Setpoint was 0.4 nN, with a tip velocity of 11.3 µm/s and a line rate of 1 Hz for 512 lines. All topographical images were processed using Gwyddion^[62]^ version 2.6.7. Second-degree polynomial row alignment and horizontal scar corrections were applied to the raw images. Mean-squared roughness (RMS) of height irregularities is reported as calculated using Gwyddion.

Using the same type of probe, force-volume measurements were performed to further extract mechanical maps from at least two 2×2 µm^2^ areas per condition, consisting of 16384 force-distance curves per 2×2 µm^2^ area. In-run of force-distance profiles were fitted with the Hertz^[63,64]^ model to estimate the Young’s modulus, with JPKSPM Data Processing version 8.1.62.

Particularly, for every force-distance curve, the entire in-run of the force measurements was considered, with the 4%-20% region of the full run analyzed, where 0% corresponds to the last point of the in-run and 100% corresponds to the starting point, preventing measuring artifacts due to the effect of the gold substrate in the mechanical characterization of the films. One of the flaws of using this method, particularly with “sharp” AFM tips, is the unknown area of contact, added to the uncertainty in the value of Poisson’s ratio, crucial in the Hertz model, which was assumed as 0.5, which for soft materials is not necessarily the case^[65,66]^, even though it can be a first approximation for highly hydrated materials. To simplify direct comparison of QCM-D shear-compliance and the AFM extracted normal Young’s modulus (*E*), normal compliance *S* values are reported as the inverse of the Young’s modulus as well, *i.e*., *S*=1/*E*. *E* distributions for all conditions are reported as histograms with a bin size of 0.33 MPa, overlapping measurements on the probed areas. Mean and standard deviation (STD) for each distribution were calculated using a Gaussian fit in OriginLab. For each condition, the average of the means is reported as *E*_avg_, along with the average of their STDs.

## RESULTS AND DISCUSSION

### Saturation of crystals was reached with 50 µg/mL Col-II and Col-I solutions

First, to determine the bulk concentration of collagen (II and I) that saturates the SAM-functionalized gold crystals, chemisorption experiments were performed with QCM-D at collagen concentrations ranging from 10 to 100 µg/mL. Col-II saturated crystals above 50 µg/mL, whereas Col-I saturated crystals at all tested concentrations (Figure S4 (A)). Shear compliance did not show dependence over the tested range of concentrations for any of the collagens (Figure S4 (B)). Based on this, 50 µg/mL was chosen as the concentration for collagen film formation.

### Collagen at neutral pH exists in a partially denaturated state

It is well established that environmental conditions, such as pH, have an impact on collagen molecular conformation, modulating the stability of the collagen triple helix^[49,51,67,68]^. To assess the conformational state of Col-II and Col-I prior to surface immobilization, CD spectra in PBS solution at pH 7 and pH 4 were acquired. Col-II presented a negative peak at ∼197 nm, with a zero-rotation point at ∼ 213 nm, and no positive maximum at ∼ 222 nm (as opposed to Col-II with perfect triple helical structure^[69]^. The negative peak at 197 nm indicates the presence of *⍺*-helices, likely originating from the individual polypeptide chains of Col-II (Figure 3 (A)). In contrast, Col-I at pH 7 and pH 4 showed clear triple-helix signatures in the CD spectrogram (Figure 3 (A)), with a negative peak at 197 nm, a zero-rotation point at 213 nm, and a positive maximum at 222 nm, consistent with previous reports^[68,70]^. However, pH 7 induced conformational changes relative to pH 4, as evidenced by a decrease in the *MRE* amplitude of both peaks at pH 7. In the case of Col-II, *R*_pn_ is lower than that of Col-I in both pH 4 and pH 7, suggesting partial denaturing of the triple helical structure, regardless of pH. While the *R*_pn_ of Col-II at pH 4 and 7 are comparable, it should be noted that the absolute values of the positive and negative peaks decreased at pH 7, suggesting a certain degree of overall disordering taking place in the switch from acidic to neutral pH. While the *R*_pn_ as a ratio of absolute peak values is in principle independent of sample concentration, a uniform flattening of the curve would not be reflected in the value, even though it is a strong indicator of collagen degradation^[71]^. In view of this, Col-I at pH 4 appears to exhibit the canonical triple-helical structure, while Col-II is found to exist in a partially degraded state. Furthermore, pH 7 seems to trigger partial degradation of both Col-I and Col-II.

**Figure 3.**
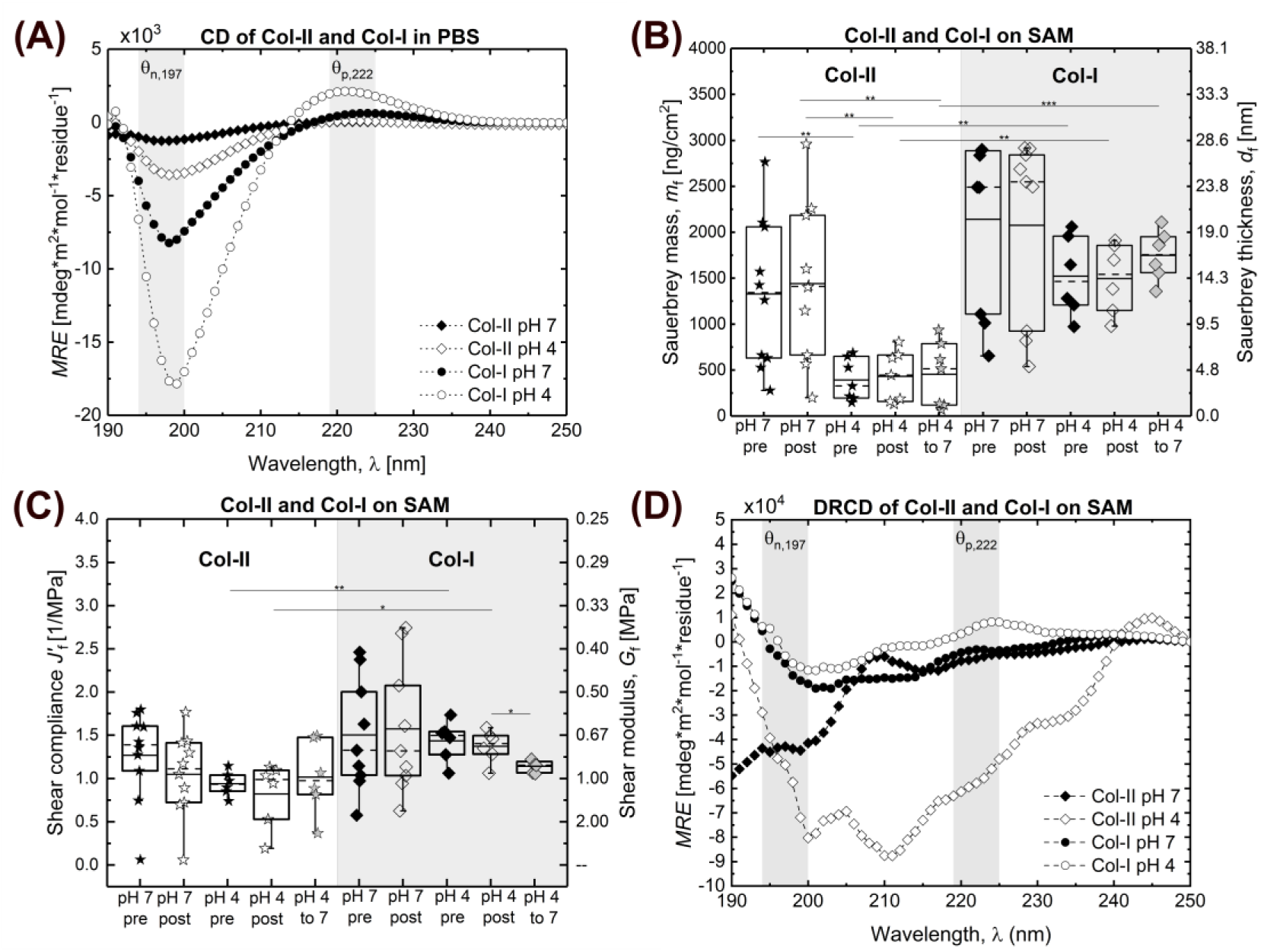
(A) Far UV CD spectra of Col-II and Col-I at pH 7 and 4 in bulk. (B) mass density and (C) thin film compliance of Col-II and Col-I immobilized on SAM before (pre) and after (post) wash with PBS at their respective pH, followed by a PBS pH 7 wash for the samples deposited at pH 4 (shown as pH 4 to 7), quantified via QCM-D. For conditions starting at pH 4, an additional wash with PBS at pH 7 is shown. Box-and-whisker plots in (C) and (D) display the mean (solid line), median (broken line), lower and upper quartiles (box) and minimum/maximum values excluding outliers with a coefficient of 1.5 (whiskers). Each data point is an independent biological replicate. Statistical differences shown are deduced from unpaired *t*-test. *p* < 0.05 is denoted as one asterisk, *p* < 0.01 as two asterisks, and *p* < 0.001 as three asterisks. (D) Far UV DRCD spectra of Col-II and Col-I deposited from PBS at pH 7 and 4, immobilized on SAM.

### Col-II film assembly was pH-dependent

As demonstrated by CD spectra (Figure 3A), pH affected the molecular conformation of Col-II and Col-I in bulk. To test whether pH-induced conformational changes affected the mass density and mechanical properties of collagen films, QCM-D was used. Collagen nanofilms were characterized by estimating their hydrated mass densities, as determined by their mass density, *m*_f_, (Figure 3(B)), and shear-dependent film compliances, 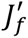 (Figure 3(C)).

Figure 3(B) shows the *m*_f_ of the pre- and post-wash of immobilized Col-II and Col-I films at pH 7 and 4, taken from at least N = 6 independent measurements. The corresponding quartiles (box), mean (line), and median (dashed line), are reported as box and whisker plots. Inferring from two-way ANOVA tests, as shown in Figure S4, collagen type and film formation pH were both variables that significantly impacted *m*_f_. No differences were observed between pre- and post-wash collagen films, indicating that the molecular content initially present in the films was retained after the PBS washing steps. Thus, the discussion for *m*_f_ focuses on the post-wash conditions. Additional *t*-tests revealed that Col-II films at pH 7 formed denser (thicker) films than those at pH 4, but this was not the case for Col-I. The mean film thicknesses of Col-II films at pH 7 were ∼ three times higher than those of films formed at pH 4. Lastly, the difference in film thickness between Col-II and Col-I formed at pH 7 was not statistically significant. This is not the case for film thicknesses at pH 4, where Col-II films were ∼ four times thinner than Col-I.

It is evident that pH impacted *m*_f_ of Coll-II films more than *m*_f_ of Col-I films. This might arise from several factors, including molecular differences (homotrimer in Col-II versus heterotrimer in Col-I), differences in the interruption of the Gly-X-Y domains, higher-order structure, and differences in isoelectric points (*I*_P_). Both Col-II and Col-I have an overall positive charge in acidic conditions (pH 4), as their *I*_P_ values are ∼ 8 and ∼ 7, respectively, resulting in a larger difference in surface charge for Col-II as compared to Col-I between the explored pH values. These factors likely led to differences in packing densities and, thus, hydration states between Col-II and Col-I films.

For films deposited at pH 4, a subtle swelling was observed when transitioning to pH 7 (pH 4 to 7 conditions in Figure 3(B)), but it was not enough to be considered statistically relevant based on *t*-test analysis.

### Col-II formed less compliant films than Col-II when deposited at pH 4

To further understand the mechanical properties and dissipative nature of the collagen films, we computed their shear-dependent thin film compliance, 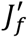. For all conditions, no significant relative changes between pre-and post-wash for 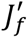 were measured. However, 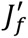 of Col-II and Col-I films at pH 4 were statistically different, as determined by a *t*-test. Interestingly, there was a significant difference in 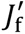 between Col-I at pH 4 post-wash and after transitioning to pH 7, showing less compliant films (with higher shear modulus) after the pH change.

The means and standard error of the means (SEMs) for the QCM-D measurements are found in Table S2. Statistical analysis results, based on *t*-tests and two-way ANOVA, for *m*_f_ and 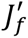 are shown in Table S3, S4, S5, and S6. The trends for the QCM-D measured properties are similar for the case of physiosorbed proteins on bare Au, although the values of adsorbed mass density are, in general, higher, as shown in Figure S5.

To gain further information on the conformation of the molecules present in the formed films, DRCD spectra of the chemisorbed films were collected. Figure 3(D) shows that most of the structural signatures were lost for both collagens when immobilized to the surfaces at acidic pH. We hypothesize that this is due to the chemical immobilization process itself disrupting the collagen molecular structure. In our previous work on fibronectin films, we observed that the immobilization strategy significantly impacts the DRCD signal^[53]^. Nevertheless, DRCD spectra of Col-I at pH 4 and pH 7 agrees with bulk CD data, as canonical Col-I peak amplitudes were more pronounced as compared to Col-II. As in bulk, peak amplitudes were also less pronounced at pH 7 as compared to pH 4, again indicating some degradation of the Col-I structure at pH 7.

The above applied photo- and acoustic-spectroscopy techniques provided conformational and nano-rheological information on the impact of pH on Col-II and Col-I film formation. To further obtain morphological and nanomechanical information of the films, atomic force microscopy (AFM) in liquid was employed, as discussed next.

### Col-II and Col-I formed continuous films with similar nanoscale morphologies across conditions

To validate that the collagen precursor films were continuous and gain insights into their morphologies, the surfaces of nanofilms fully immersed in liquid (PBS at pH 7) were imaged using AFM in QI mode, as shown in Figure 4. Col-II and Col-I, regardless of the deposition pH, formed continuous films with globular features. Additionally, larger 5×5 µm^2^ area scans are presented in Figure S6.

**Figure 4.**
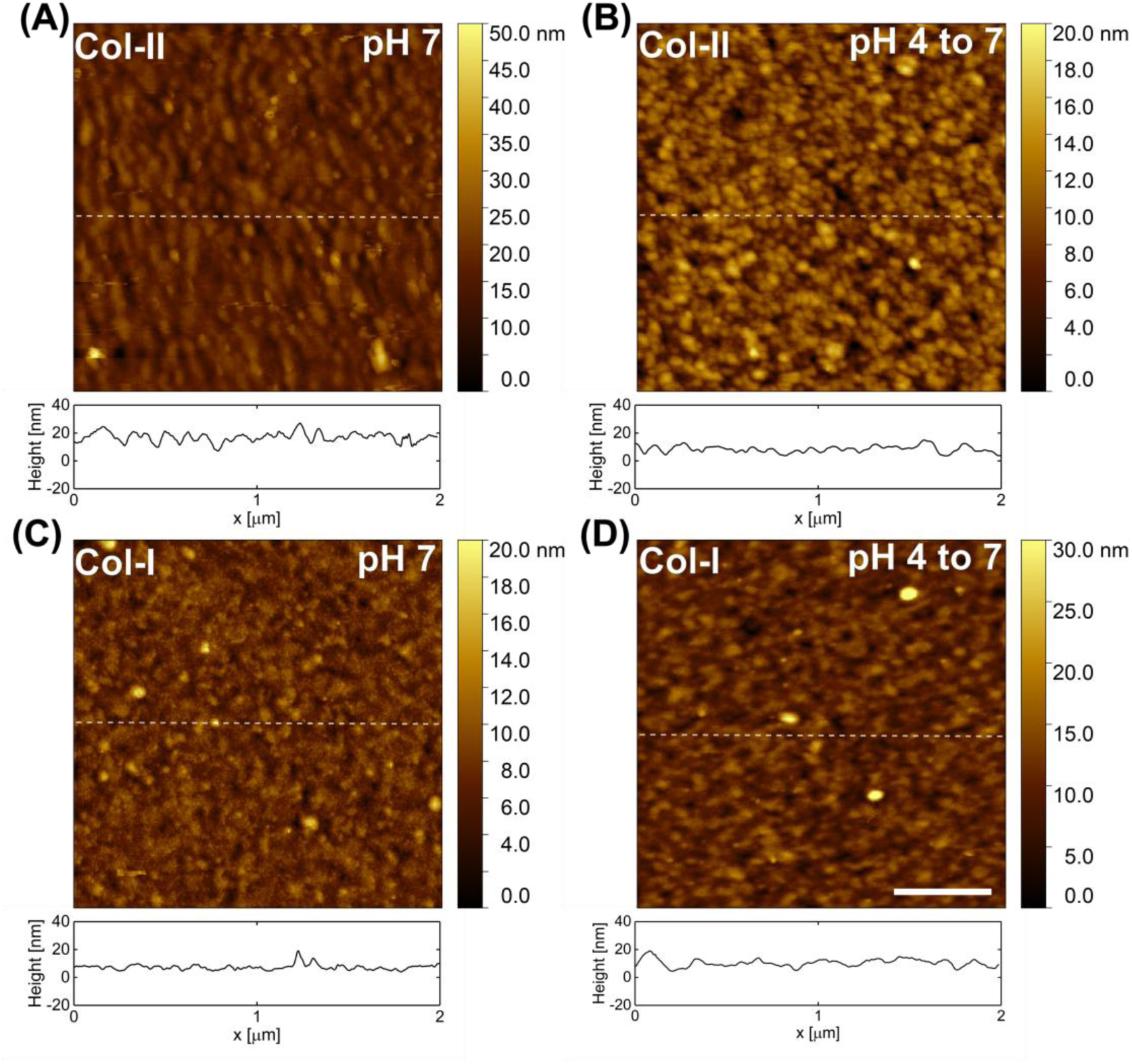
Representative AFM height maps of: (A) Col-II pH 7, (B) Col-II pH 4, (C) Col-I pH 7, (D) Col-I pH 4. All conditions were measured in a PBS liquid environment at pH 7. Representative cross sections of each map are shown below them. Scale bar in (D) applies to all micrographs of this figure and is 500 nm. At least two independent biological replicates were measured for each condition, showing similar results.

**Figure 5.**
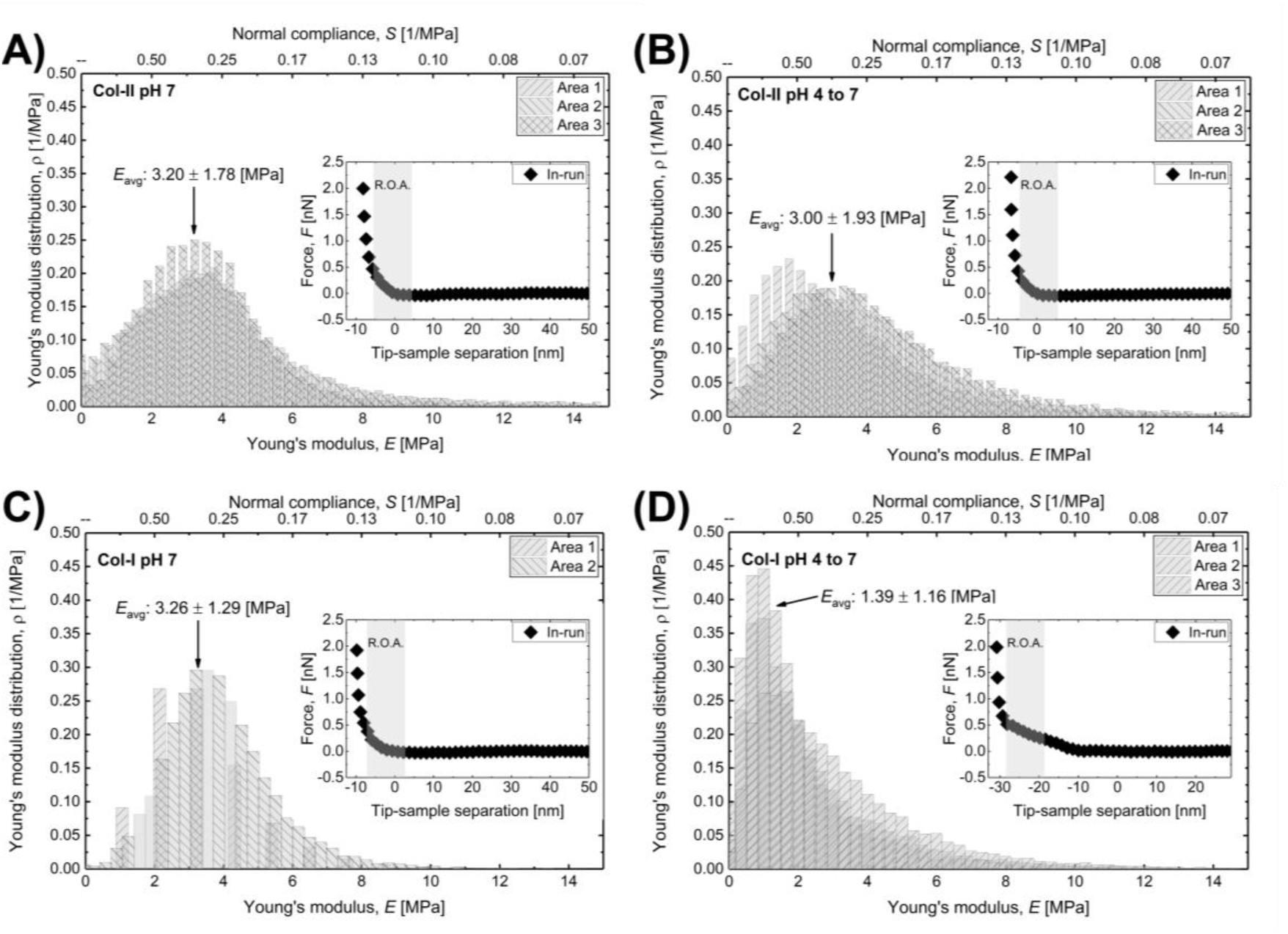
AFM Young’s moduli distributions of (A) Col-II pH 7, (B) Col-II deposited at pH 4 and measured at pH 7 (pH 4 to 7), (C) Col-I pH 7, (D) Col-I pH 4 to 7. All conditions were measured in a PBS liquid environment at pH 7. A secondary horizontal axis (top) reports normal compliance, *S*, as the inverse of the Young’s moduli (*S*=1/*E*). Insets are representative force-distance curves for each condition, with their region of analysis (R.O.A.) used to extract *E*. *E*_avg_ indicates the average of the means obtained for each individual distribution as shown in Figure S7. Each “Area” histogram represents an independent technical replicate.

2×2 µm^2^ micrographs of Col-II films deposited at pH 7 show surfaces with an RMS roughness of 4 nm, while those deposited from pH 4 had an RMS roughness of 3 nm. Col-II formed at pH 7 presented slightly more elongated structures that appeared to have a preferred orientation along the slow axis of the AFM scan. These elongated structures can also be seen in the 5×5 µm^2^ area scans, in Figure S6(C). Col-I films deposited at pH 7 had an RMS roughness of 3 nm, while those deposited at pH 4 had an RMS roughness of 2 nm. Similar morphologies and RMS roughness values of physiosorbed Col-II films on gold under similar experimental conditions have been reported in the literature^[29]^.

In environments with relatively high ionic strength, it has been shown that salt ions screen intermolecular interactions between Col-II triple helices, negatively impacting collagen microfibril formation^[52,72]^. This effect begins to appear at physiological ionic strengths (∼150-200 mM), a range that includes the ionic strength of our PBS buffer (∼160 mM). This can be a reason for the lack of large microfibril structures in the AFM micrographs shown (in addition to the low concentrations of Col-II and Col-I of 0.05 mg/ml used in this study). To further understand the normal nanomechanics of Coll-II and Col-I films, another independent set of samples was characterized by force-volume mapping using AFM, as discussed next.

### Col-II films formed stiffer than Col-I films when deposited at pH 4

Next, we tested if the trends and magnitudes observed by nano-rheological properties measured via QCM-D (shear-compliance) held for nanoindentation via AFM.

Nanomechanical characterization based on force-volume mapping performed on each of the precursor collagen films revealed differences in Young’s modulus (or compliance) between Col-II and Col-I films deposited from pH 4, but not between Col-II and Col-I films deposited at pH 7. Furthermore, the distribution of the Young’s moduli for Col-II deposited at pH 4 showed a normal distribution, with a mean and standard deviation (STD) of 3.00 ± 1.93 MPa, while for Col-I deposited at pH 4, the distribution of Young’s moduli resembles more a right-skewed normal distribution with a mean and STD of 1.39 ± 1.16 MPa. That is, Col-II films deposited at pH 4 showed a mean of *E* of ∼ double than that of *E* for Col-I. For films formed from pH 7, the difference is barely noticeable, having means and STDs of 3.20 ± 1.78 MPa and 3.26 ± 1.29 MPa for Col-II and Col-I, respectively. Notice that the mean and STD reported are the average of the mean and STD of the independent areas for each condition, which were all independently fitted with Gaussian functions, as shown in Figure S7. All collagen films, however, showed some degree of hysteresis between the in- and out-runs, as shown in the representative force-distance curves in Figure S8. The response of collagen films to the interaction with the AFM probe (which had a maximum ∼ 2 nN, for a radius of curvature ∼ 20 nm) was not purely elastic, as the recovery (if any) was not immediate. This is not surprising, as collagen is known to have a viscoelastic response under nanoindentation^[73]^. These findings, however, show that this characteristic of collagen is most likely present even when the triple-helical structure is presumably lost to a large extent (in agreement with DRCD). Young’s moduli values of these nanofilms also are consistent with previously reported values for collagen gels, obtained from nanoindentation^[74]^.

Connecting the nano-rheological properties obtained from QCM-D and the nano-mechanics for AFM, if the collagen films are considered isotropic, homogeneous, and linearly elastic, the Young’s modulus *E* should be approximately three times the shear modulus *G*_f_, which is in good agreement with the elasticity measurements of Col-II. However, this is not the case for Col-I, suggesting a deviation from an isotropic, homogeneous, and linearly elastic assumption (Table S7).

### Col-II acted as a scaffold for dilute synovial fluid (dSF) and recombinant lubricin (rLub), while Col-I adsorbed and retained rLub only

To assess the capability of the Col-II and Col-I films to scaffold the supramolecular assembly of dSF and rLub films, QCM-D was used. Collagen films formed in the QCM-D system, as described above, were further exposed to dSF and rLub. The *m*_f_ of the adsorbed and retained dSF and rLub on the collagen films was quantified and is shown in Figure 6.

**Figure 6.**
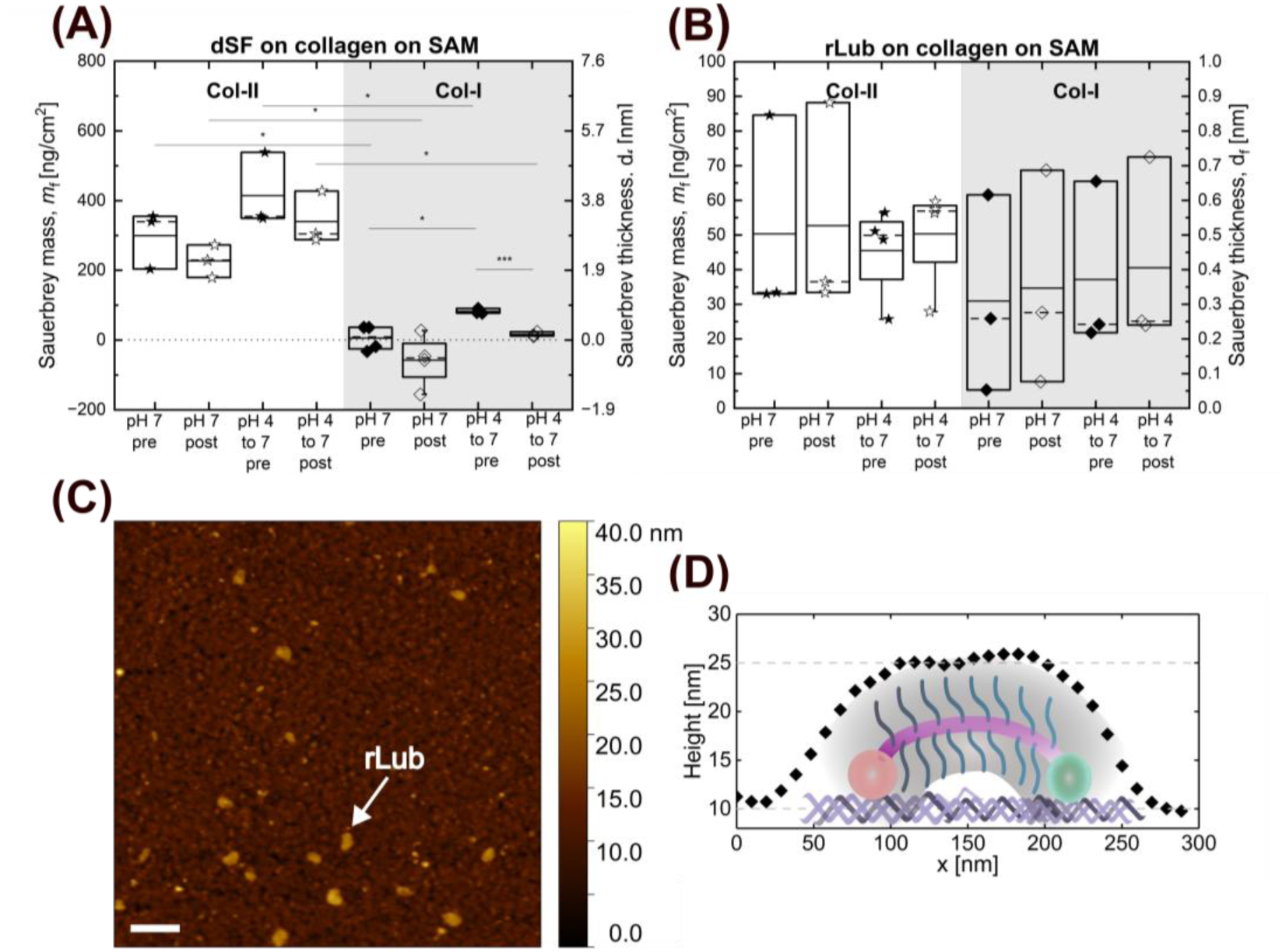
QCM-D computed *m*_f_ of (A) dSF and (B) rLub adsorbed and retained layers on collagen precursor films. Note that the vertical axes are on different scales for (A) and (B). Box-and-whisker plots in (A) and (B) display the mean (solid line), median (broken line), lower and upper quartiles (box) and minimum/maximum values excluding outliers with a coefficient of 1.5 (whiskers). Each data point is an independent biological replicate. Statistical differences shown are deduced from unpaired *t*-test. *p* < 0.05 is denoted as one asterisk, *p* < 0.01 as two asterisks, and *p* < 0.001 as three asterisks. (C) AFM height image (contact mode) of rLub on a Col-I film. Scale bar is 500 nm. (D) Average height profile of 5 rLub molecules of (C), to visualize a possible conformation of rLub molecules attached to the surface.

No significant difference in dSF adsorption on Col-II deposited at pH 7 vs. pH 4 was measured. The same observations hold true for post-wash *m*_f_ values. Unlike Col-II films, Col-I at pH 7 showed *m*_f_ values close to zero or negative. These measurements are reported as “little or no adsorption” in Table S8. One plausible interpretation of the negative *m*_f_ values is that dSF desorbs Col-I molecules without adsorbing. An alternative view is that *m*_f_ is decreasing due to the desorption of Col-I molecules, with lighter dSF constituents replacing Col-I at a rate that thins the overall participating layers. This can be attributed to the complex interactions between the plethora of macromolecular constituents present in dSF. A third interpretation is that surface bound molecules, likely from the previous blocking step (BSA), were displaced, due to the Vroman effect^[56]^, by lighter dSF constituents with higher affinity for Col-I. It is important to note that the blocking molecule, BSA, is a major constituent of SF ^[11]^.

On Col-I films deposited at pH 4, unlike those at pH 7, the *m*_f_ of dSF was positive. After the wash, this amount significantly decreased, indicating a loss of most of the adsorbed mass. Both, adsorbed and retained dSF films on Col-I at pH 4 to 7 were, however, significantly smaller than the *m*_f_ for dSF on either Col-II condition.

Overall, the type of collagen mediates the scaffolding of dSF, as reflected by dSF *m*_f_ (or, analogously, *d*_f_ thickness of dSF films formed), supported by *t*-tests and two-way ANOVA, as shown in Tables S9 and S10. This finding alone might be crucial for understanding the decrease in mechanical performance of compromised synovial joints, considering that the Col-II to Col-I ratio decreases in pathologies such as osteoarthritis and rheumatoid arthritis^[47]^. Furthermore, this finding can assist in the design of biomaterials for cartilage repair or bioactive coatings for orthopedic implants that are directly exposed to SF.

Next, we explored the role of Col-II and Col-I in scaffolding rLub, a molecule identified as a key regulator of friction and synovial joint mechanics^[13,75]^ present in SF, and proposed as a therapy for pathological synovial joints^[22]^. A key requirement of boundary lubricants is the ability to adsorb and remain attached to the surface, as validated for rLub via QCM-D.

### Col-II and Col-I adsorbed and retained similar amounts of rLub molecules for all conditions

We quantified the adsorption of rLub directly on Col-II and Col-I films formed at pH 7 and 4, as shown in Figure 6(B). For all conditions tested, no statistically significant differences in their *m*_f_ were observed. This contrasts with dSF *m*_f_ findings, suggesting that the assembly of dSF films on the precursor films was not driven solely by lubricin, but by intermolecular synergies among SF components, as has been reported for HA and Lub^[18]^, HA and PLs^[4]^ and HA with Aggrecans^[76]^. *m*_f_ values for rLub are summarized in Table S8.

Based on the conversion from *m*_f_ to *d*_f_ (Sauerbrey thickness) used in this study, rLub films across conditions should have thicknesses less than a nanometer if they were continuous and homogeneous. However, this is not realistic, as a single hydrated lubricin molecule is expected to have a cross-sectional diameter of ∼10 nm. Our results suggest that the rLub molecules did not form continuous, homogeneous films, but rather, rLub decorated collagen surfaces discretely. Indeed, supporting the interpretation of a discretely decorated collagen film, AFM imaging of rLub on Col-I is shown in Figure 6(C). This has been observed and reported for rLub on Col-II via AFM as well^[29]^, with the same protein concentrations used in our study.

Furthermore, the average cross-section of five different rLub molecules on Col-I is presented in Figure 6(D), along with a schematic of a possible conformation of the rLub monomers. This is, assuming that the rLub monomers are either lying flat on the Col-I surface, or forming loops, assuming they attached through their C- and N-termini to the underlying collagens. Another possibility is that of rLub multimers interconnecting through their N-termini, remained bound to the Col-I surface via their C-termini only, as has been proposed previously for lubricin binding to fibronectin^[11]^. The exact way in which lubricin decorates AC-ECM surfaces to provide excellent lubrication remains an open question. Results of *t*-test analysis of *m*_f_ of the adsorbed rLub films are presented in Table S9, and two-way ANOVA results in Table S11.

### dSF disrupted the mechanical integrity of Col-I films

Lastly, to gain insights into the compliance (which incorporates contributions of the QCM-D dissipation channel to the modeling, while *m*_f_ does not) of the dSF and rLub on Col-II and Col-I stack system, shear compliance 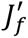 of the stack, was quantified. Results of the post-wash conditions are shown in Figure 7. Pre- and post-wash 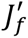 values for the stacked systems are presented in Table S12. Results of t-test statistical analysis of the stacked systems is reported in Table S13.

**Figure 7.**
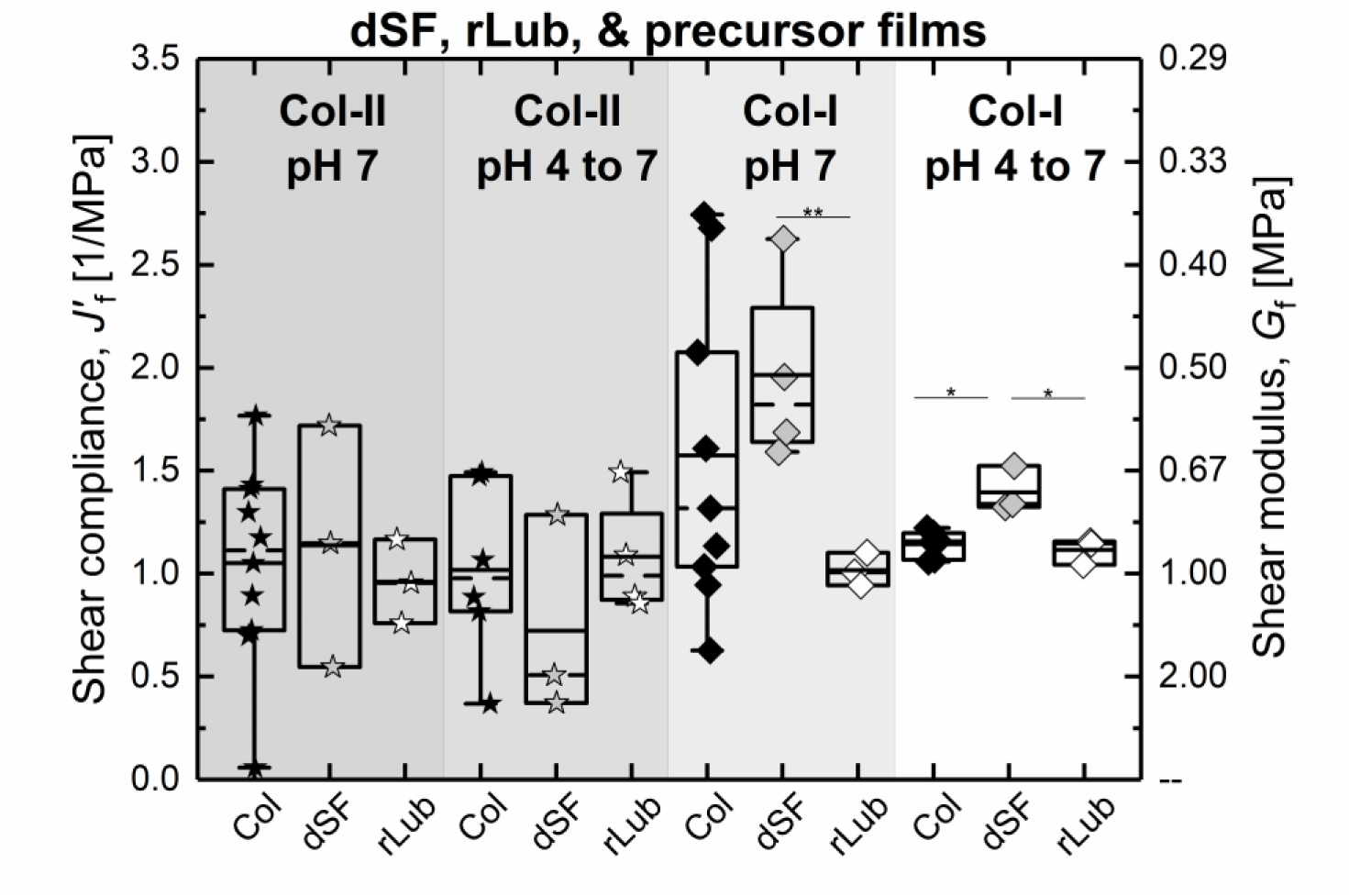
Shear-film compliance of post-wash wash conditions for Col-II and Col-I films with adsorbed dSF or rLub. Box-and-whisker plot displays the mean (solid line), median (broken line), lower and upper quartiles (box) and minimum/maximum values excluding outliers with a coefficient of 1.5 (whiskers). Each data point is an independent biological replicate. Statistical differences shown are deduced from unpaired *t*-test. *p* < 0.05 is denoted as one asterisk, *p* < 0.01 as two asterisks, and *p* < 0.001 as three asterisks.

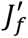 post-wash values for dSF and collagen films revealed no statistically significant differences for any of the reported values, as shown in Figure 7(A). When comparing the stacked films of collagen and the adlayers (dSF or rLub) with their respective conditions of collagen alone, significant difference between the shear compliance of Col-I pH 4 to 7 and dSF + Col-I pH 4 to 7 was revealed, with a more compliant system when synovial fluid components are adsorbed to the precursor film than when only rLub is adsorbed. For films with collagen and rLub, there was no statistically significant difference in any condition compared with their respective measurements with collagen alone. Interestingly, statistically significant differences were observed between the stacks of Col-I and dSF, and the stack of Col-I and rLub, for both, Col-I formed at pH 7 and at pH 4. This indicates that the stack of dSF and Col-I was more compliant than that of rLub and Col-I at both formation pHs. Considering that there was no statistical difference between the stacked rLub and collagen systems, while there was with respect to the dSF and collagen systems (for Col-I), the “little to no” adsorption of dSF in Col-I systems still affected the mechanical properties of the film. This was even more apparent for the condition of Col-I pH 4 to 7, in which the collagen film was thinner than when deposited from pH 7. Film compliance is influenced by the molecular network’s cross-linking, water content, and protein density, among other factors. This suggests that dSF disrupted the Col-I films, as shown in Figure 7, thereby affecting their mechanical integrity. This effect was not evidenced on Col-II films, where dSF components were likely able to form denser layers.

Control experiments. Control experiments previously reported elsewhere^[53]^ by our group consisted of quantifying mf of dSF on bare Au surfaces, Au surfaces with SAM, and Au surfaces with SAM and a BSA blocking step, to gain insights into the role of collagen films in adsorbing and retaining dSF. In all these control conditions, mf of dSF was significantly higher than mf of dSF on the collagen films that reported in this work, suggesting that the dSF adsorption on collagen films was specific to collagens and not to the underlying bare gold or SAM surfaces. The adsorption of BSA alone on Au and collagen films is shown in Figure S9. Adsorption of R-LUB on bare gold and SAM with glutaraldehyde in shown in Figure S10.

## CONCLUSIONS

In this work, molecular films of Col-II and Col-I were chemically immobilized to self-assembled monolayers on gold, at two different initial pHs, 4 and 7. DRCD, QCM-D, and AFM showed differences in collagen conformations, nanomorphology, and mechanical properties among the films, with thicker, more compliant, and more structured films forming at neutral pH. Furthermore, collagen films were exposed to synovial fluid (dSF) and recombinant lubricin (rLub), to gain insight into their ability to regulate the supramolecular assembly and retention of dSF and rLub films. Our results show that, when exposed to SF, differences in adsorption of up to 100% between Col-II and Col-I were detected. This was not the case for rLub, which adsorbed forming a discrete, non-continuous films with similar mass densities across all tested conditions, suggesting that lubricin alone does not drive full SF film assembly in our cartilage models. Our findings provide insight into the mechanisms underlying the loss of mechanical performance in injured cartilage, driven by Col-II loss and alterations in the Col-II:Col-I ratios. Thus, the collagen isoform is a parameter that should not be overlooked when designing functional biological biomaterials intended for the AC-ECM (synthetic cartilage implants). Future efforts will focus on elucidating the tribological performance of these bio-interfaces.

## Supporting information

None

## ACKNOWLEDGMENTS

R.C.A.E., D.R.J.P., and U.H. acknowledge funding from the National Science Foundation (NSF) CAREER Award through NSF-CMMI-2239665 awarded to R.C.A.E. R.C.A.E., D.R.J.P., S.T.A., N.M., K.C., S.W., M.J.P., H.R., and L.J.B. acknowledge NSF LEAP HI funding through NSF-CMMI-2245367 awarded to L.J.B., M.J.P, H.R., and R.C.A.E. We acknowledge funding from NSF-CREST: Center for Cellular and Biomolecular Machines through NSF-HRD-1547848. D.R.J.P. and R.C.A.E. thank Dr. Mourad Sadqi for technical support and assistance during this project. K.D. acknowledges funding from the Natural Sciences and Engineering Research Council of Canada (NSERC) Discovery Grants Program through RGPIN-2023–03607. D.R.J.P. would like to thank Sixin Zhai and Alex Mears, from the MBSE graduate group at UC Merced, for giving great feedback during the write-up of this manuscript.

## SUPPLEMENTARY INFORMATION

Figure S1. Friction coefficient of rLub and PBS lubricating cartilage on glass surfaces at varying sliding speeds.

Figure S2. Representative QCM-D plots of change in frequency and dissipation for the experiments where dSF was flowed on the collagen films.

Figure S3. Representative QCM-D plots of change in frequency and dissipation for the experiments where rLub was flowed on the collagen film.

Figure S4. QCM-D measurements of immobilized collagen films on SAM functionalized gold-coated crystals at pH 7 as a function of bulk concentration of Col-II and Col-I between 10 – 100 µg/ml.

Figure S5. QCM-D measurements of collagen films physiosorbed on bare Au.

Figure S6. 5×5 µm^2^ AFM height maps of immobilized collagen films.

Figure S7. AFM Young’s moduli distributions and Gaussian fit on different 2×2 μm^2^ areas of the immobilized collagen films.

Figure S8. Force-runs obtained with AFM force-volume mapping the immobilized collagen films.

Figure S9. Sauerbrey mass, *m*_f_, of BSA adsorbed onto various surfaces at pH 7.

Figure S10. Sauerbrey mass, *m*_f_, of rLub adsorbed on bare gold and gold with SAM and glutaraldehyde at pH 7.

Table S1. Parameters used for DRCD modelling.

Table S2. Summary of QCM-D determined collagen film properties before (pre) and after (post) PBS washes on SAM.

Table S3. Two-way ANOVA test results for Sauerbrey mass of collagen films.

Table S4. *t*-test results or Sauerbrey mass of collagen films.

Table S5. *t*-test results for shear dependent compliance of collagen films

Table S6. Two-way ANOVA test results for shear dependent compliance of collagen films.

Table S7. Comparison of mechanical properties measured by AFM and QCM.

Table S8. Summary of QCM-D determined dSF and rLub film Sauerbrey masses before (pre) and after (post) PBS washes on collagen films.

Table S9. *t*-test results for Sauerbrey mass of rLub and dSF films, presented in Figure 6 and the discussion of the main text.

Table S10. Two-way ANOVA test results for Sauerbrey mass of dSF on collagen films.

Table S11. Two-way ANOVA test results for Sauerbrey mass of rLub on collagen films.

Table S12. Summary of shear-dependent compliance determined for dSF and rLub and their respective collagen films.

Table S13. *t*-test results for shear dependent compliance of the stacked rLub and dSF films with collagen.

## REFERENCES

[1] T. A. Schmidt, N. S. Gastelum, Q. T. Nguyen, B. L. Schumacher, R. L. Sah, Arthritis Rheum. 2007, 56, 882.

[2] J. P. Gleghorn, A. R. C. Jones, C. R. Flannery, L. J. Bonassar, Journal of Orthopaedic Research 2009, 27, 771.

[3] W. Lin, J. Klein, Acc. Mater. Res. 2022, 3, 213.

[4] K. Sun, M. W. Rutland, R. M. Espinosa-Marzal, APPLIED SCIENCES AND ENGINEERING Lipid self-assembly dependence on hyaluronic acid size reveals biolubrication and osteoarthritic degeneration mechanisms, Vol. 12, 2026.

[5] S. Jahn, J. Seror, J. Klein, Annu. Rev. Biomed. Eng. 2016, 18, 235.

[6] B. Zappone, G. W. Greene, E. Oroudjev, G. D. Jay, J. N. Israelachvili, Langmuir 2008, 24, 1495.

[7] R. C. Andresen Eguiluz, S. G. Cook, M. Tan, C. N. Brown, N. J. Pacifici, M. S. Samak, L. J. Bonassar, D. Putnam, D. Gourdon, Front. Bioeng. Biotechnol. 2017, 5.

[8] I. S. Bayer, Lubricants 2018, 6.

[9] A. R. C. Jones, J. P. Gleghorn, C. E. Hughes, L. J. Fitz, R. Zollner, S. D. Wainwright, B. Caterson, E. A. Morris, L. J. Bonassar, C. R. Flannery, Journal of Orthopaedic Research 2007, 25, 283.

[10] G. D. Jay, D. A. Harris, C.-J. Cha, Boundary lubrication by lubricin is mediated by O-linked β(1-3)Gal-GalNAc oligosaccharides, Vol. 18, Kluwer Academic Publishers, 2001.

[11] R. C. Andresen Eguiluz, S. G. Cook, C. N. Brown, F. Wu, N. J. Pacifici, L. J. Bonassar, D. Gourdon, Biomacromolecules 2015, 16, 2884.

[12] B. J. Wu, Q. Y. Deng, Y. X. Leng, C. M. Wang, N. Huang, Surf. Coat. Technol. 2017, 320, 320.

[13] A. S. Mann, A. M. Smith, J. O. Saltzherr, A. Gopinath, R. C. Andresen Eguiluz, Colloids Surf. B Biointerfaces 2022, 213.

[14] I. Onu, R. Gherghel, I. Nacu, F. D. Cojocaru, L. Verestiuc, D. V. Matei, D. Cascaval, I. L. Serban, D. A. Iordan, A. Tucaliuc, A. I. Galaction, Biomedicines 2024, 12.

[15] D. A. Swann, R. B. Hendren, E. L. Radin, S. L. Sotman, Arthritis Rheum. 1981, 24, 22.

[16] S. G. Cook, Y. Guan, N. J. Pacifici, C. N. Brown, E. Czako, M. S. Samak, L. J. Bonassar, D. Gourdon, Langmuir 2019.

[17] X. Banquy, D. W. Lee, S. Das, J. Hogan, J. N. Israelachvili, Adv. Funct. Mater. 2014, 24, 3152.

[18] S. Das, X. Banquy, B. Zappone, G. W. Greene, G. D. Jay, J. N. Israelachvili, Biomacromolecules 2013, 14, 1669.

[19] G. W. Greene, X. Banquy, D. W. Lee, D. D. Lowrey, J. Yu, J. N. Israelachvili, .

[20] K. Jerczynski, D. A. Pham, A. Kucherova, A. Filippini, J. Robert, J. Ulanski, F. Moldovan, X. Banquy, J. Pietrasik, Chemical Engineering Journal 2025, 169341.

[21] B. Zappone, M. Ruths, G. W. Greene, G. D. Jay, J. N. Israelachvili, Biophys. J. 2007, 92, 1693.

[22] M. J. Paszek, H. L. Reesink, RECOMBINANT LUBRICINS, AND COMPOSITIONS AND METHODS FOR USING THE SAME, USA, 2025.

[23] C. R. Shurer, Y. Wang, E. Feeney, S. E. Head, V. X. Zhang, J. Su, Z. Cheng, M. A. Stark, L. J. Bonassar, H. L. Reesink, M. J. Paszek, Biotechnol. Bioeng. 2019, 116, 1292.

[24] M. J. Colville, L.-T. Huang, S. Schmidt, K. Chen, K. Vishwanath, J. Su, R. M. Williams, L. J. Bonassar, H. L. Reesink, M. J. Paszek, 2024.

[25] S. J. Womack, E. J. Secor, S. R. Nelissen, A. G. Fortin, A. M. Schiller, M. J. Colville, L. J. Bonassar, M. J. Paszek, H. L. Reesink, Sci. Rep. 2026.

[26] M. Tschaikowsky, S. Brander, V. Barth, R. Thomann, B. Rolauffs, B. N. Balzer, T. Hugel, Acta Biomater. 2022, 146, 274.

[27] M. Pei, C. Yu, M. Qu, Expression of collagen type I, II and III in loose body of osteoarthritis, Vol. 5, 2000.

[28] D. P. Chang, F. Guilak, G. D. Jay, S. Zauscher, J. Biomech. 2014, 47, 659.

[29] H. Yuan, L. L. E. Mears, X. Liu, W. Qi, R. Su, M. Valtiner, Colloids Surf. B Biointerfaces 2022, 220.

[30] S. A. Flowers, A. Zieba, J. Örnros, C. Jin, O. Rolfson, L. I. Björkman, T. Eisler, S. Kalamajski, M. Kamali-Moghaddam, N. G. Karlsson, Sci. Rep. 2017, 7.

[31] J. S. Pieper, P. M. Van Der Kraan, T. Hafmans, J. Kamp, P. Buma, J. L. C. Van Susante, W. B. Van Den Berg, J. H. Veerkamp, T. H. Van Kuppevelt, Crosslinked type II collagen matrices: preparation, characterization, and potential for cartilage engineering, Vol. 23, 2002.

[32] V. Irawan, T. C. Sung, A. Higuchi, T. Ikoma, Collagen Scaffolds in Cartilage Tissue Engineering and Relevant Approaches for Future Development, Vol. 15, Korean Tissue Engineering and Regenerative Medicine Society, 2018, pp. 673–697.

[33] P. Buma, J. S. Pieper, T. Van Tienen, J. L. C. Van Susante, P. M. Van Der Kraan, J. H. Veerkamp, W. B. Van Den Berg, R. P. H. Veth, T. H. Van Kuppevelt, Biomaterials 2003, 24, 3255.

[34] P. Mafi, S. Hindocha, R. Mafi, W. S. Khan, Evaluation of Biological Protein-Based Collagen Scaffolds in Cartilage and Musculoskeletal Tissue Engineering-A Systematic Review of the Literature, Vol. 7, 2012.

[35] H. Zhou, Z. Zhang, Y. Mu, L. Ma, X. Hu, B. Liu, D. A. Wang, Advanced Materials 2025.

[36] H. Yuan, H. W. Cheng, L. L. E. Mears, R. Huang, R. Su, W. Qi, Z. He, M. Valtiner, Langmuir 2021, 37, 13810.

[37] M. L. Smith, D. Gourdon, W. C. Little, K. E. Kubow, R. A. Eguiluz, S. Luna-Morris, V. Vogel, PLoS Biol. 2007, 5, 2243.

[38] E. Klotzsch, M. L. Smith, K. E. Kubow, S. Muntwyler, W. C. Little, F. Beyeler, D. Gourdon, B. J. Nelson, V. Vogel, Fibronectin forms the most extensible biological fibers displaying switchable force-exposed cryptic binding sites, 2009.

[39] W. C. Little, R. Schwartlander, M. L. Smith, D. Gourdon, V. Vogel, Nano Lett. 2009, 9, 4158.

[40] K. Wang, R. C. Andresen Eguiluz, F. Wu, B. R. Seo, C. Fischbach, D. Gourdon, Biomaterials 2015, 54, 63.

[41] E. M. Chandler, M. P. Saunders, C. J. Yoon, D. Gourdon, C. Fischbach, Phys. Biol. 2011, 8.

[42] E. M. Chandler, B. R. Seo, J. P. Califano, R. C. Andresen Eguiluz, J. S. Lee, C. J. Yoon, D. T. Tims, J. X. Wang, L. Cheng, S. Mohanan, M. R. Buckley, I. Cohen, A. Y. Nikitin, R. M. Williams, D. Gourdon, C. A. Reinhart-King, C. Fischbach, Proc. Natl. Acad. Sci. U. S. A. 2012, 109.

[43] B. Ri Seo, X. Chen, L. Ling, Y. Hye Song, A. A. Shimpi, S. Choi, J. Gonzalez, J. Sapudom, K. Wang, R. Carlos Andresen Eguiluz, D. Gourdon, V. B. Shenoy, C. Fischbach, .

[44] J. Kim, L. J. Bonassar, J. Biomed. Mater. Res. A 2023, 111, 478.

[45] A. Sorushanova, L. M. Delgado, Z. Wu, N. Shologu, A. Kshirsagar, R. Raghunath, A. M. Mullen, Y. Bayon, A. Pandit, M. Raghunath, D. I. Zeugolis, The Collagen Suprafamily: From Biosynthesis to Advanced Biomaterial Development, Vol. 31, Wiley-VCH Verlag, 2019.

[46] P. Cui, T. Shao, W. Liu, M. Li, M. Yu, W. Zhao, Y. Song, Y. Ding, J. Liu, Advanced review on type II collagen and peptide: preparation, functional activities and food industry application, Vol. 64, Taylor and Francis Ltd., 2024, pp. 11302–11319.

[47] Z. Ouyang, L. Dong, F. Yao, K. Wang, Y. Chen, S. Li, R. Zhou, Y. Zhao, W. Hu, Cartilage-Related Collagens in Osteoarthritis and Rheumatoid Arthritis: From Pathogenesis to Therapeutics, Vol. 24, Multidisciplinary Digital Publishing Institute (MDPI), 2023.

[48] M. J. Buehler, Nature designs tough collagen: Explaining the nanostructure of collagen fibrils, 2006.

[49] Y. Wang, L. Zhang, W. Liao, Z. Tong, F. Yuan, L. Mao, J. Liu, Y. Gao, Food Hydrocoll. 2023, 137.

[50] F. Jiang, H. Hörber, J. Howard, D. J. Müller, J. Struct. Biol. 2004, 148, 268.

[51] E. Bianchi, G. Conio, A. Ciferri, D. Puett, L. Rajagh, J. Biol. Chem. 1967, 242, 1361.

[52] S. Morozova, M. Muthukumar, Journal of Chemical Physics 2018, 149.

[53] S. T. Ahmed, D. R. Jaramillo Pinto, L. Vitkova, U. Honey, W. Flores, K. L. Lunny, K. A. Cutter, Y. Wen, K. De France, R. C. Andresen Eguiluz, Colloids Surf. B Biointerfaces 2026, 258, 115224.

[54] M. J. Colville, L.-T. Huang, S. Schmidt, K. Chen, K. Vishwanath, J. Su, R. M. Williams, L. J. Bonassar, H. L. Reesink, M. J. Paszek, Recombinant manufacturing of multispecies biolubricants, 2024.

[55] J. P. Gleghorn, L. J. Bonassar, J. Biomech. 2008, 41, 1910.

[56] S. L. Hirsh, D. R. McKenzie, N. J. Nosworthy, J. A. Denman, O. U. Sezerman, M. M. M. Bilek, Colloids Surf. B Biointerfaces 2013, 103, 395.

[57] B. Pardi, S. T. Ahmed, S. J. Flores, W. Flores, J.-M. Friedt, L. L. E. Mears, B. Y. Soto, R. C. A. Eguiluz, J. Open Source Softw. 2024, 9, 6831.

[58] J. Vörös, Biophys. J. 2004, 87, 553.

[59] B. Du, D. Johannsmann, Langmuir 2004, 20, 2809.

[60] A. Savitzky, M. J. E, Smoothing and Differentiation of Data by Simplified Least Squares Procedures, Vol. 40, 1951.

[61] E. Khare, C.-H. Yu, C. Gonzalez Obeso, M. Milazzo, D. L. Kaplan, M. J. Buehler, M. J. B. Designed, M. J. B. Performed, M. J. B. Contributed, 2022, 119, 2209524119.

[62] D. Neèas, P. Klapetek, Gwyddion: An open-source software for SPM data analysis, Vol. 10, 2012, pp. 181–188.

[63] V. Herrn Heinrich Hertz, lieber die Berührung fester elastischer Körper.

[64] E. Dintwa, E. Tijskens, H. Ramon, Granul. Matter 2008, 10, 209.

[65] A. M. Smith, D. G. Inocencio, B. M. Pardi, A. Gopinath, R. C. Andresen Eguiluz, ACS Appl. Polym. Mater. 2024, 6.

[66] K. K. Dwivedi, P. Lakhani, S. Kumar, N. Kumar, R. Soc. Open Sci. 2022, 9.

[67] C. Y. Liu, M. Matsusaki, M. Akashi, Polym. J. 2015, 47, 391.

[68] T. Ikoma, H. Kobayashi, J. Tanaka, D. Walsh, S. Mann, Int. J. Biol. Macromol. 2003, 32, 199.

[69] Y. Wang, S. Yang, L. Zhang, F. Yuan, L. Mao, J. Liu, Y. Gao, Food Chem. 2023, 406.

[70] H. Tian, Z. Ren, L. Shi, G. Hao, J. Chen, W. Weng, Process Biochemistry 2021, 108, 153.

[71] M. Sun, X. Wei, H. Wang, C. Xu, B. Wei, J. Zhang, L. He, Y. Xu, S. Li, Food Bioproc. Tech. 2020, 13, 367.

[72] K. G. Wilcox, G. M. Kemerer, S. Morozova, Journal of Chemical Physics 2023, 158.

[73] M. Asgari, E. Mirzarazi, R. J. Benavides, Y. M. Efremov, R. D. Frisina, H. Vali, H. D. Espinosa, Acta Biomater. 2026, 209, 466.

[74] Y. Olguín, M. Selva, D. Benavente, N. Orellana, I. Montenegro, A. Madrid, D. Jaramillo-Pinto, M. C. Otero, T. P. Corrales, C. A. Acevedo, Pharmaceutics 2023, 15, 2760.

[75] G. D. Jay, J. R. Torres, M. L. Warman, M. C. Laderer, K. S. Breuer, The role of lubricin in the mechanical behavior of synovial fluid, 2007.

[76] J. Seror, Y. Merkher, N. Kampf, L. Collinson, A. J. Day, A. Maroudas, J. Klein, Biomacromolecules 2012, 13, 3823.

