## Supplementary material for "Functional Differentiation of Type II and Type I Collagen Articular Models in Synovial Fluid Film Formation and Recombinant Lubricin Retention": None

[1]Materials and Biomaterials Science and Engineering Graduate Program, University of California, Merced, Merced, USA, [2]Department of Chemical and Materials Engineering, School of Engineering, University of California, Merced, Merced, USA, [3]Department of Chemical Engineering, Smith Engineering, Queen's University, Kingston, ON, Canada, [4]Sibley School of Mechanical and Aerospace Engineering, Cornell University, Ithaca, NY, USA, [5]Department of Biomedical Engineering, University of California, Davis, Davis, CA, USA, [6]Department of Surgical and Radiological Sciences, School of Veterinary Medicine, University of California, Davis, Davis, CA, USA, [7]Meinig School of Biomedical Engineering, Cornell University, Ithaca, NY, USA, [8]Health Sciences Research Institute, University of California, Merced, Merced, USA

**Corresponding author**

*

**Figure S1.** Friction coefficient of rLub and PBS lubricating cartilage on glass surfaces at varying sliding speeds.

**Figure S2.** Representative QCM-D plots of change in frequency and dissipation for the experiments where dSF was flowed on the collagen films.

**Figure S3.** Representative QCM-D plots of change in frequency and dissipation for the experiments where rLub was flowed on the collagen film.

**Figure S4.** QCM-D measurements of immobilized collagen films on SAM functionalized gold-coated crystals at pH 7 as a function of bulk concentration of Col-II and Col-I between 10 – 100 μg/ml.

**Figure S5.** QCM-D measurements of collagen films physiosorbed on bare Au.

**Figure S6.** 5x5 $\mu m^2$ AFM height maps of immobilized collagen films.

**Figure S7.** AFM Young's moduli distributions and Gaussian fit on different 2x2 $\mu m^2$ areas of the immobilized collagen films.

**Figure S8.** Force-runs obtained with AFM force-volume mapping the immobilized collagen films.

**Figure S9.** Sauerbrey mass, $m_f$, of BSA adsorbed onto various surfaces at pH 7.

**Figure S10.** Sauerbrey mass, $m_f$, of rLub adsorbed on bare gold and gold with SAM and glutaraldehyde at pH 7.

**Table S1.** Parameters used for DRCD modelling.

**Table S2.** Summary of QCM-D determined collagen film properties before (pre) and after (post) PBS washes on SAM.

**Table S3.** Two-way ANOVA test results for Sauerbrey mass of collagen films.

**Table S4.** *t*-test results or Sauerbrey mass of collagen films.

**Table S5.** *t*-test results for shear dependent compliance of collagen films

**Table S6.** Two-way ANOVA test results for shear dependent compliance of collagen films.

**Table S7.** Comparison of mechanical properties measured by AFM and QCM.

**Table S8.** Summary of QCM-D determined dSF and rLub film Sauerbrey masses before (pre) and after (post) PBS washes on collagen films.

**Table S9.** *t*-test results for Sauerbrey mass of rLub and dSF films, presented in Figure 6 and the discussion of the main text.

**Table S10.** Two-way ANOVA test results for Sauerbrey mass of dSF on collagen films.

**Table S11.** Two-way ANOVA test results for Sauerbrey mass of rLub on collagen films.

**Table S12.** Summary of shear-dependent compliance determined for dSF and rLub and their respective collagen films.

**Table S13.** *t*-test results for shear dependent compliance of the stacked rLub and dSF films with collagen.

**Lubricating properties of rLub.** A custom pin-on-plate tribometer was used to quantify the lubricating performance of the utilized recombinant lubricin on cartilage against glass, in a strain control mode. The neo-natal bovine cartilage (88 Meat Commodities LLC, USA) piece with 6 mm in diameter and 2 mm of thickness is submerged in a rLub solution of 5 mg/ml. Each independent sample was compressed to 30% axial strain and then allowed to stress-relax for one hour, followed by sliding. The friction coefficients were calculated as the ratio between the average shear load and the average normal load at the end of the sliding when friction had reached an equilibrium, measured at sliding speeds of 0.1 mm/s, 1 mm/s, and 10 mm/s. The same process was done for control with PBS only, as shown below. Figure SX shows that for the measured system, rLub in PBS solution reduced the friction coefficient by half an order of magnitude with respect to PBS alone.

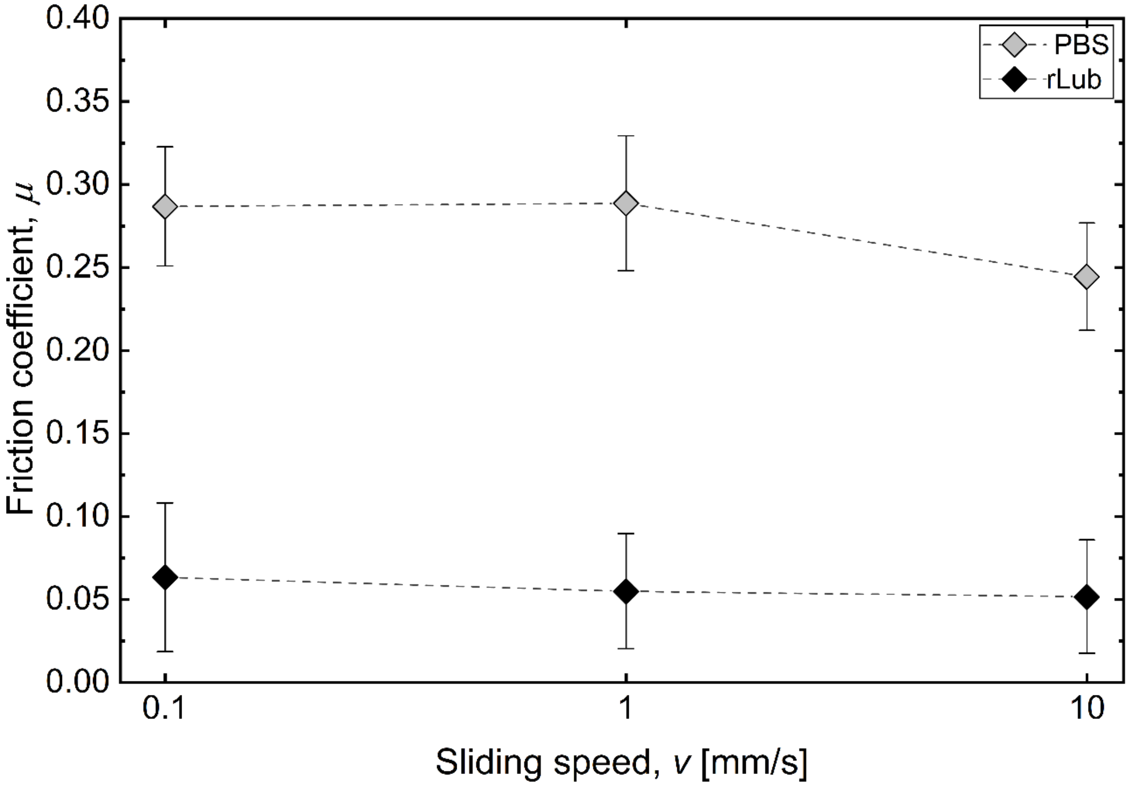


**Figure S1.** Friction coefficient of rLub (N = 3 independent measurements) and PBS (N = 4 independent measurements) lubricating cartilage on glass surfaces at varying sliding speed.

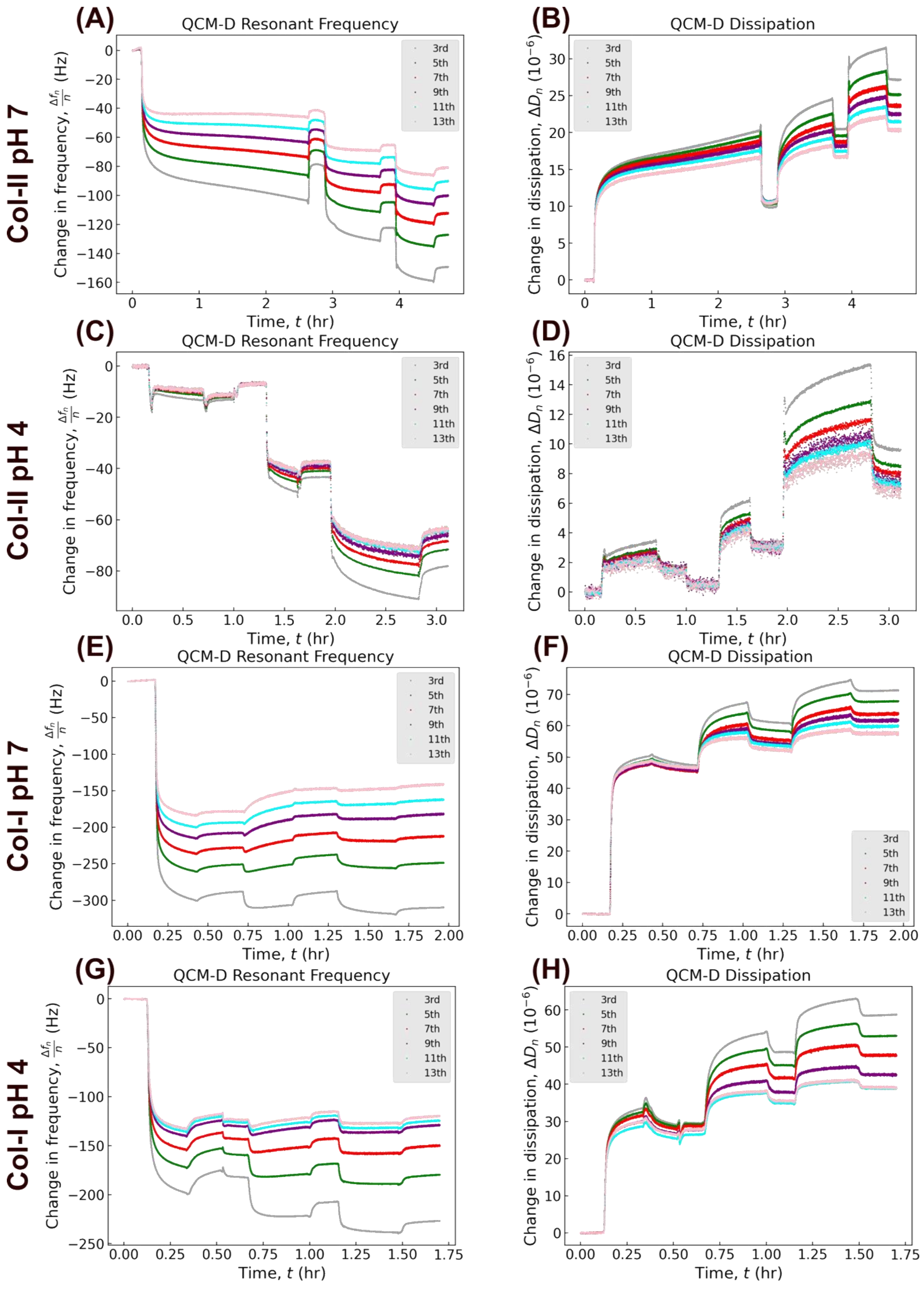


**Figure S2.** Representative QCM-D plots of change in frequency and dissipation for the experiments where dSF was flowed on the collagen films, with a prior BSA blocking step.

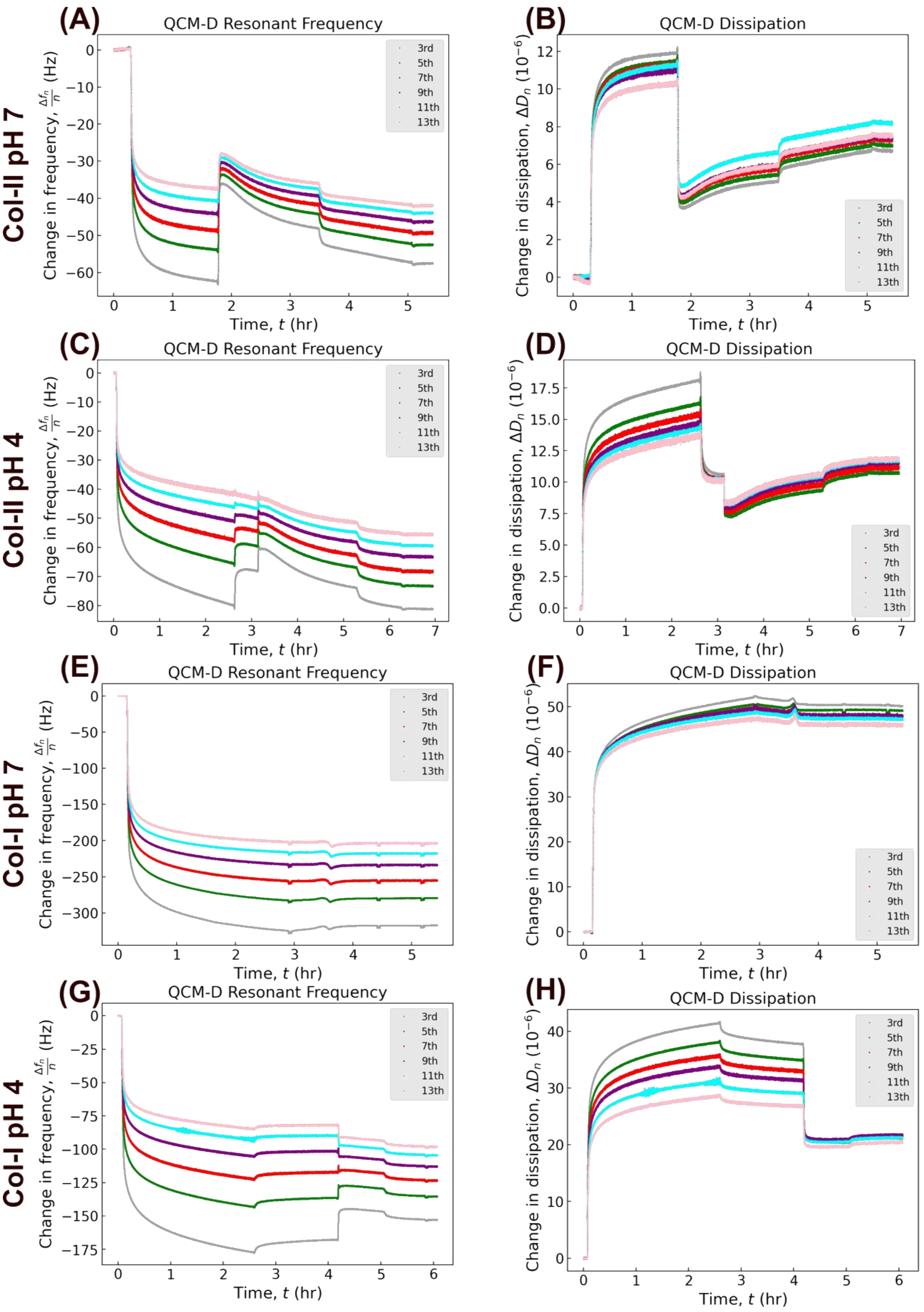


**Figure S3.** Representative QCM-D plots of change in frequency and dissipation for the experiments where rLub was flowed on the collagen film.

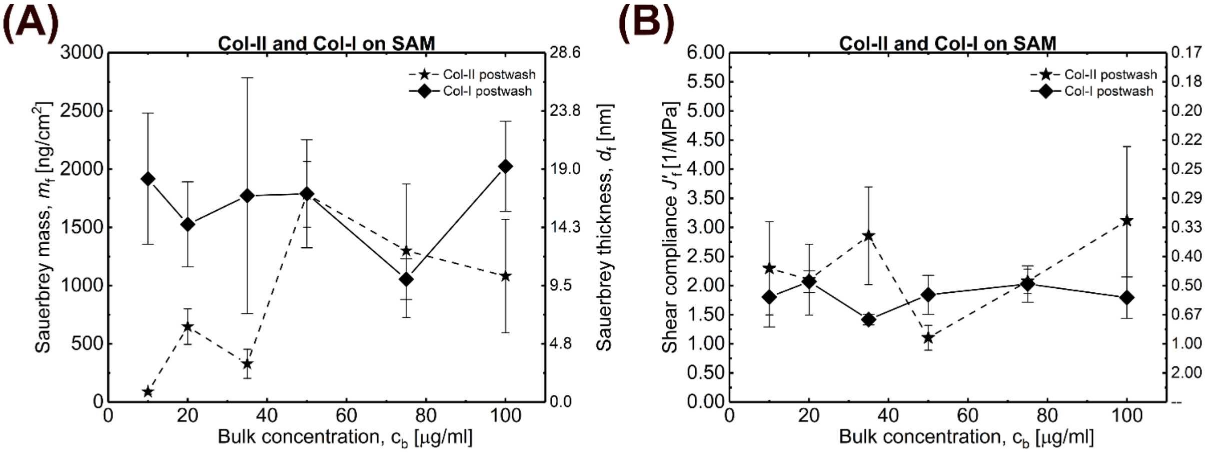


**Figure S4.** Sauerbrey mass (A) and shear compliance (B) of collagen films on SAM functionalized gold-coated crystals at pH 7 as a function of bulk concentration of Col-II and I between 10 – 100 μg/ml.

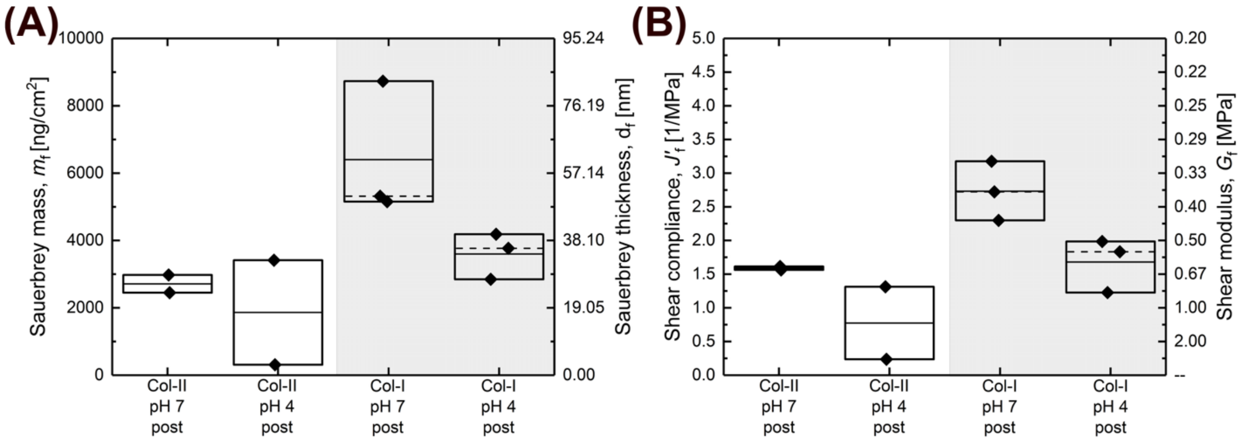


**Figure S5.** Sauerbrey mass (A) and shear compliance (B) and Far UV DRCD spectra of collagen physiosorbed on bare Au (no SAM).

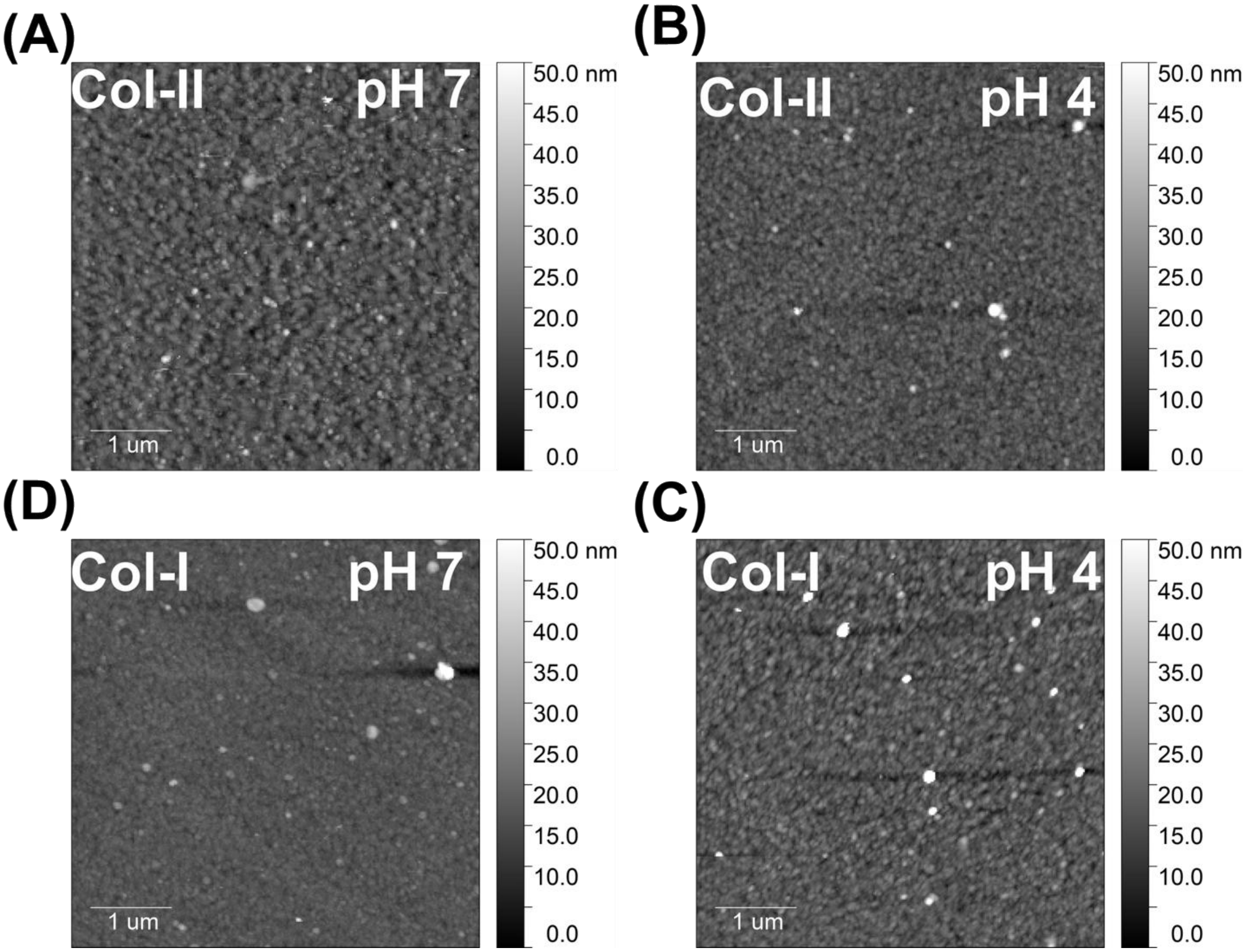


**Figure S6.** 5x5 μm$^2$ AFM height maps of (A) Col-II pH 7, (B) Col-II pH 4, (C) Col-I pH 7, (D) Col-I pH 4. All conditions were measured in a PBS liquid environment at pH 7.

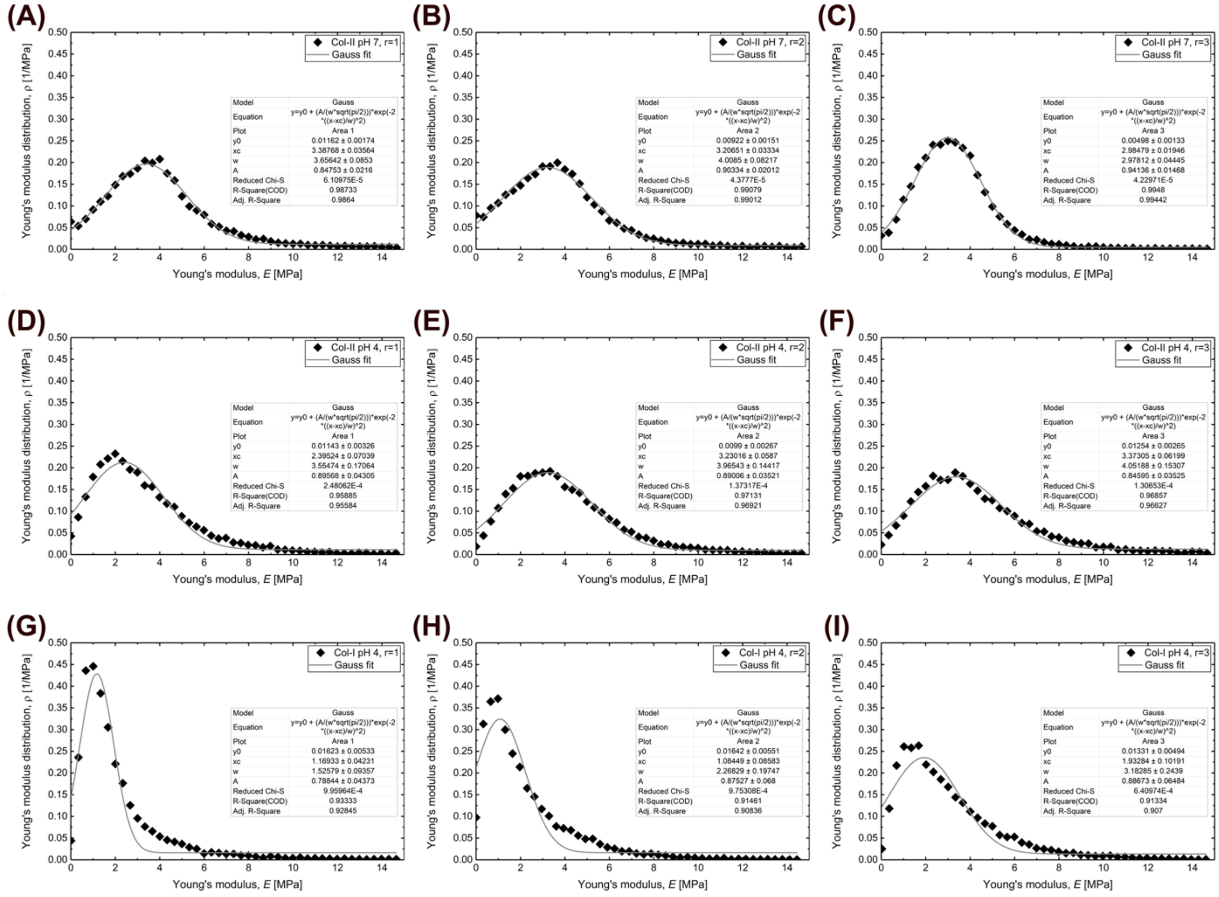


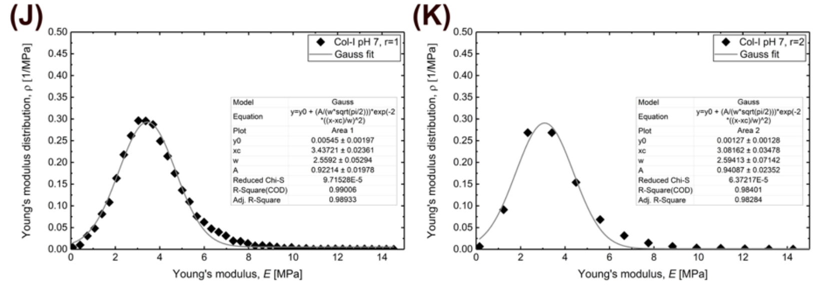


(L)

| Protein - pH | Col-II pH 7 | Col-II pH 4 | Col-I pH 7 | Col-I pH 4 |
|---|---|---|---|---|
| Young's modulus [MPa] | $3.20 \pm 1.78$ | $3.00 \pm 1.93$ | $3.26 \pm 1.29$ | $1.39 \pm 1.16$ |

**Figure S7.** AFM Young's modulus distributions and Gaussian fit on different 2x2 $\mu m^2$ areas for (A, B, C) Col-II pH 7, (D, E, F) Col-II pH 3, (G, H, I) Col-I pH 4, and (J, K) Col-I pH 7. (L) shows the averages of the means for each condition, as well as the averages of their standard deviation.

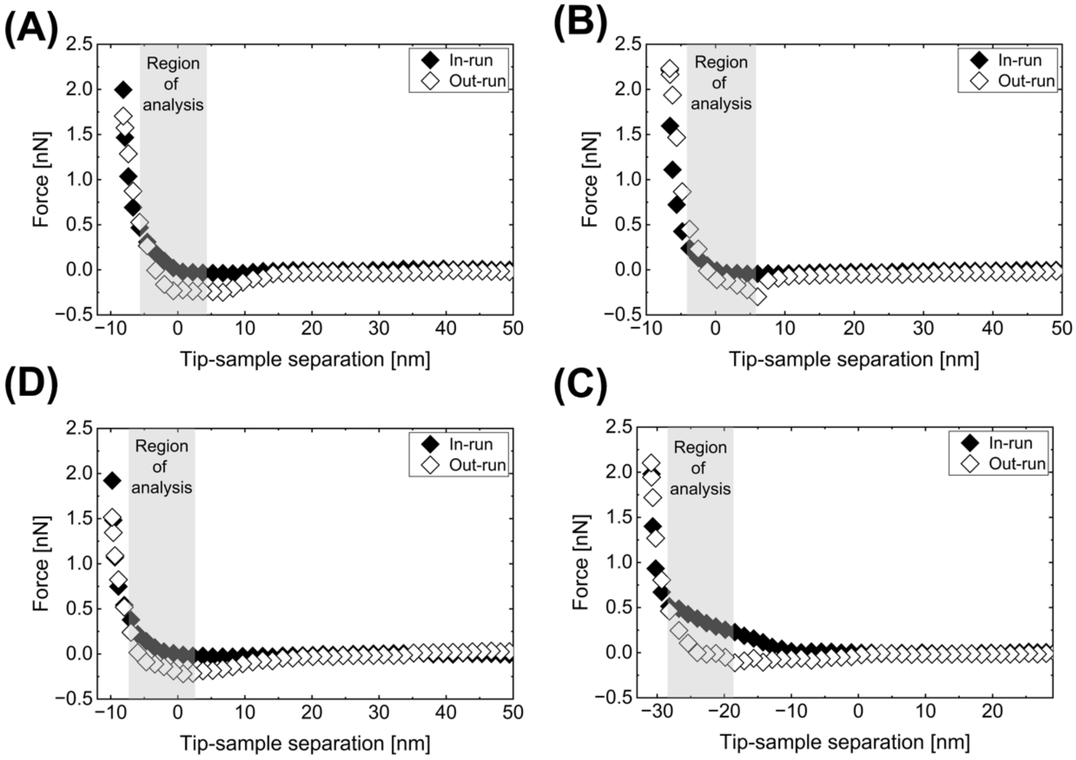


**Figure S8.** Force-runs obtained with AFM force-volume mapping of (A) Col-II pH 7, (B) Col-II pH 4, (C) Col-I pH 7, (D) Col-I pH 4. All conditions were measured in a PBS liquid environment at pH 7.

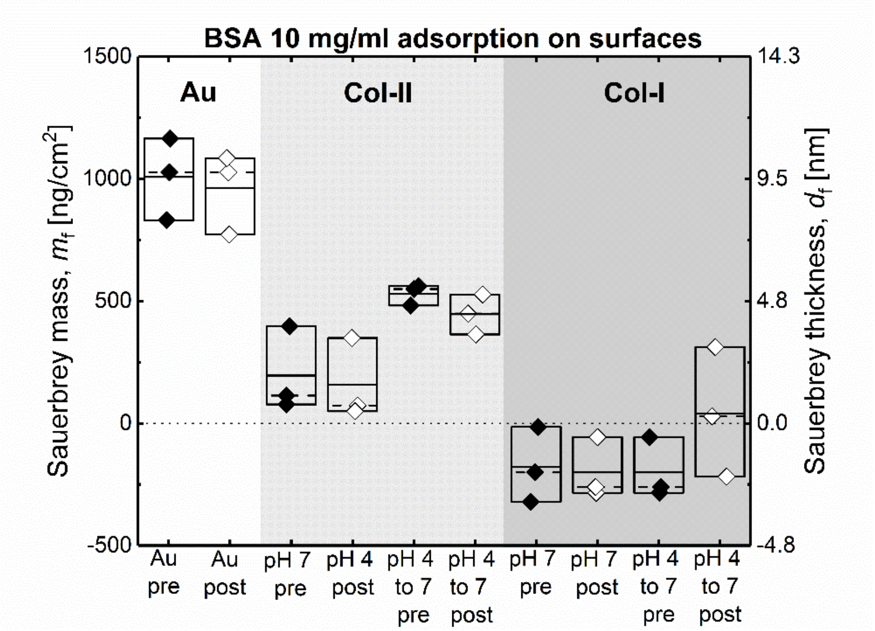


**Figure S9.** Sauerbrey mass, $m_f$, of BSA adsorbed onto various surfaces, all in pH 7. The Au condition is clean, bare Au (and has been previously reported by our group[1]). The Col-II and Col-I surfaces are immobilized on SAM.

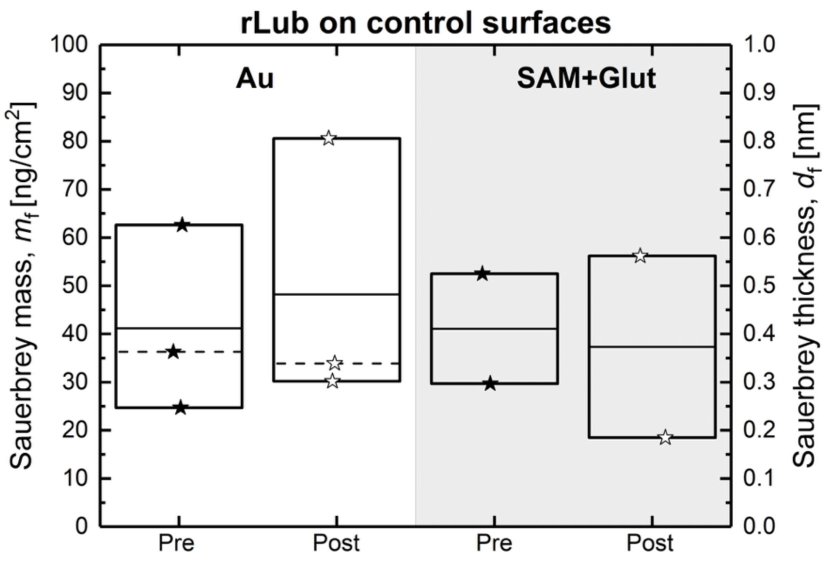


**Figure S10.** Sauerbrey mass, $m_f$, of rLub adsorbed onto bare Au and functionalized Au with SAM and glutaraldehyde, all at pH 7.

**Table S1.** Parameters used for DRCD modeling, applied in eq. 6 of the main text.

| | $M_w$ [g/mol] | $m_f$ [ng/cm$^2$] | $N_r$ |
|---|---|---|---|
| Short SAM+ GLU+Col-II pH7 | 300000 | 1441 | 3000 |
| Short SAM+ GLU+Col-I pH7 | 300000 | 1800 | 3000 |
| Short SAM+GLU+Col-II pH4 | 300000 | 454 | 3000 |
| Short SAM+GLU+Col-I pH4 | 300000 | 1748 | 3000 |

**Table S2.** Summary of QCM-D determined collagen film properties before (pre) and after (post) PBS washes on SAM.

| Protein - pH | Pre-wash $m_f$ [ng/cm$^2$] | Post-wash $m_f$ [ng/cm$^2$] | pH 4 to 7 $m_f$ [ng/cm$^2$] | Pre-wash $J_f'$ [1/MPa] | Post-wash $J_f'$ [1/MPa] | pH 4 to 7 $J_f'$ [1/MPa] |
|---|---|---|---|---|---|---|
| Col-II - pH 7 | 1328 ± 257 | 1441 ± 270 | - | 1.27 ± 0.17 | 1.05 ± 0.15 | - |
| Col-II - pH 4 | 394 ± 86 | 433 ± 104 | 454±133 | 0.94 ± 0.06 | 0.82 ± 0.15 | 1.02 ± 0.17 |
| Col-I – pH 7 | 2142 ± 311 | 1800 ± 500 | - | 1.50 ± 0.22 | 1.57 ± 0.26 | - |
| Col-I – pH 4 | 1521 ± 178 | 1496 ± 158 | 1748 ± 277 | 1.44 ± 0.10 | 1.37 ± 0.08 | 1.14 ± 0.03 |

**Table S3.** Two-way ANOVA test results for Sauerbrey mass of collagen films, presented in Figure 3 (B) of the main text.

*Overall ANOVA*

| | DF | Sum of Squares | Mean Square | F Value | P Value |
|---|---|---|---|---|---|
| CollagenType | 1 | 6.49633E6 | 6.49633E6 | 11.39235 | 0.00211 |
| FormationpH | 1 | 3.56431E6 | 3.56431E6 | 6.25058 | 0.01832 |
| Model | 2 | 1.01771E7 | 5.08857E6 | 8.92361 | 9.54674E-4 |
| Error | 29 | 1.65369E7 | 570236.24068 | -- | -- |
| Corrected Total | 31 | 2.6714E7 | -- | -- | -- |

At the 0.05 level, the population means of **CollagenType** are **significantly** different.
At the 0.05 level, the population means of **FormationpH** are **significantly** different.

**Table S4.** *t*-test results or Sauerbrey mass of collagen films, presented in Figure 3 (B) of the main text.

| Group 1 | Group 2 | Mean 1 | Mean 2 | SEM 1 | SEM 2 | n1 | n2 | t-statistic | df | p-value | Result |
|---|---|---|---|---|---|---|---|---|---|---|---|
| Col-II prewash in pH 7 | Col-II postwash in pH 7 | 1328.837 | 1440.666 | 257.2158 | 269.8518 | 10 | 10 | -0.3 | 17.96 | 0.7676 | Not Significant (p > 0.0 |
| Col-I prewash in pH 7 | Col-I postwash in pH 7 | 2141.728 | 2076.901 | 311.0886 | 334.0161 | 9 | 9 | 0.142 | 15.92 | 0.8888 | Not Significant (p > 0.0 |
| Col-II prewash in pH 4 | Col-II postwash in pH 4 | 393.696 | 433.007 | 85.62478 | 104.8645 | 7 | 7 | -0.2904 | 11.54 | 0.7767 | Not Significant (p > 0.0 |
| Col-II postwash in pH 4 | Col-II 4 to 7 | 433.007 | 454.2419 | 104.8645 | 132.3688 | 7 | 7 | -0.1257 | 11.4 | 0.9021 | Not Significant (p > 0.0 |
| Col-I postwash in pH 4 | Col-I prewash in pH 4 | 1496.282 | 1521.339 | 157.593 | 178.2759 | 6 | 6 | -0.1053 | 9.85 | 0.9182 | Not Significant (p > 0.0 |
| Col-I postwash in pH 4 | Col-I 4 to 7 | 1496.282 | 1748.272 | 157.593 | 113.027 | 6 | 6 | -1.2994 | 9.07 | 0.2259 | Not Significant (p > 0.0 |
| Col-II prewash in pH 7 | Col-I prewash in pH 7 | 1328.837 | 2141.728 | 257.2158 | 311.0886 | 10 | 9 | -2.0138 | 16.02 | 0.0611 | Not Significant (p > 0.0 |
| Col-II postwash in pH 7 | Col-I postwash in pH 7 | 1440.666 | 2076.901 | 269.8518 | 334.0161 | 10 | 9 | -1.4817 | 15.85 | 0.158 | Not Significant (p > 0.0 |
| Col-II prewash in pH 4 | Col-I prewash in pH 4 | 393.696 | 1521.339 | 85.62478 | 178.2759 | 7 | 6 | -5.7017 | 7.25 | 0.0006 | Significant (p ≤ 0.05) |
| Col-II postwash in pH 4 | Col-I postwash in pH 4 | 433.007 | 1496.282 | 104.8645 | 157.593 | 7 | 6 | -5.6171 | 8.95 | 0.0003 | Significant (p ≤ 0.05) |
| Col-II postwash in pH 7 | Col-II 4 to 7 | 1440.666 | 454.2419 | 269.8518 | 132.3688 | 10 | 7 | 3.2819 | 12.75 | 0.0061 | Significant (p ≤ 0.05) |
| Col-I postwash in pH 7 | Col-I 4 to 7 | 2076.901 | 1748.272 | 334.0161 | 113.027 | 9 | 6 | 0.932 | 9.73 | 0.3739 | Not Significant (p > 0.0 |
| Col-II 4 to 7 | Col-I 4 to 7 | 454.2419 | 1748.272 | 132.3688 | 113.027 | 7 | 6 | -7.4344 | 10.95 | 0 | Significant (p ≤ 0.05) |
| Col-II postwash in pH 7 | Col-II postwash in pH 4 | 1440.666 | 433.007 | 269.8518 | 104.8645 | 10 | 7 | 3.4806 | 11.53 | 0.0048 | Significant (p ≤ 0.05) |
| Col-I postwash in pH 7 | Col-I postwash in pH 4 | 2076.901 | 1496.282 | 334.0161 | 157.593 | 9 | 6 | 1.5721 | 11.08 | 0.144 | Not Significant (p > 0.0 |
| Col-I prewash in pH 7 | Col-I prewash in pH 4 | 2141.728 | 1521.339 | 311.0886 | 178.2759 | 9 | 6 | 1.7303 | 12.04 | 0.1091 | Not Significant (p > 0.0 |
| Col-II prewash in pH 7 | Col-II prewash in pH 4 | 1328.837 | 393.696 | 257.2158 | 85.62478 | 10 | 7 | 3.4495 | 10.9 | 0.0055 | Significant (p ≤ 0.05) |

.

**Table S5** *t*-test results for shear dependent compliance of collagen films, presented in Figure 3 (C) of the main text.

| Group 1 | Group 2 | Mean 1 | Mean 2 | SEM 1 | SEM 2 | n1 | n2 | t-statistic | df | p-value | Result |
|---|---|---|---|---|---|---|---|---|---|---|---|
| Col-II prewash in pH4 | Col-II postwash in pH4 | 0.94156 | 0.82182 | 0.05866 | 0.1544 | 6 | 6 | 0.725 | 6.41 | 0.4941 | Not Significant (p > 0.0 |
| Col-I prewash in pH4 | Col-I postwash in pH4 | 1.4354 | 1.37234 | 0.09557 | 0.07582 | 6 | 6 | 0.5169 | 9.51 | 0.617 | Not Significant (p > 0.0 |
| Col-I prewash in pH7 | Col-I postwash in pH7 | 1.5033 | 1.57403 | 0.21906 | 0.2551 | 9 | 9 | -0.2104 | 15.64 | 0.8361 | Not Significant (p > 0.0 |
| Col-II prewash in pH7 | Col-II postwash in pH7 | 1.27048 | 1.05074 | 0.16772 | 0.15298 | 10 | 10 | 0.968 | 17.85 | 0.346 | Not Significant (p > 0.0 |
| Col-II prewash in pH4 | Col-I prewash in pH4 | 0.94156 | 1.4354 | 0.05866 | 0.09557 | 6 | 6 | -4.4039 | 8.3 | 0.0021 | Significant (p ≤ 0.05) |
| Col-II postwash in pH4 | Col-I postwash in pH4 | 0.82182 | 1.37234 | 0.1544 | 0.07582 | 6 | 6 | -3.2005 | 7.28 | 0.0143 | Significant (p ≤ 0.05) |
| Col-I prewash in pH7 | Col-II prewash in pH7 | 1.5033 | 1.27048 | 0.21906 | 0.16772 | 9 | 10 | 0.8439 | 15.42 | 0.4116 | Not Significant (p > 0.0 |
| Col-I postwash in pH7 | Col-II postwash in pH7 | 1.57403 | 1.05074 | 0.2551 | 0.15298 | 9 | 10 | 1.7592 | 13.26 | 0.1016 | Not Significant (p > 0.0 |
| Col-I prewash in pH4 | Col-I prewash in pH7 | 1.4354 | 1.5033 | 0.09557 | 0.21906 | 6 | 9 | -0.2841 | 10.71 | 0.7817 | Not Significant (p > 0.0 |
| Col-I postwash in pH4 | Col-I postwash in pH7 | 1.37234 | 1.57403 | 0.07582 | 0.2551 | 6 | 9 | -0.7579 | 9.36 | 0.4672 | Not Significant (p > 0.0 |
| Col-II prewash in pH4 | Col-II prewash in pH7 | 0.94156 | 1.27048 | 0.05866 | 0.16772 | 6 | 10 | -1.8512 | 11.04 | 0.0911 | Not Significant (p > 0.0 |
| Col-II postwash in pH4 | Col-II postwash in pH7 | 0.82182 | 1.05074 | 0.1544 | 0.15298 | 6 | 10 | -1.0532 | 12.79 | 0.3117 | Not Significant (p > 0.0 |
| Col-II 4 to 7 | Col-I 4 to 7 | 1.01763 | 1.14287 | 0.1747 | 0.0275 | 6 | 6 | -0.7082 | 5.25 | 0.5091 | Not Significant (p > 0.0 |
| Col-II postwash in pH4 | Col-II 4 to 7 | 0.82182 | 1.01763 | 0.1544 | 0.1747 | 6 | 6 | -0.8398 | 9.85 | 0.4209 | Not Significant (p > 0.0 |
| Col-I postwash in pH4 | Col-I 4 to 7 | 1.37234 | 1.14287 | 0.07582 | 0.0275 | 6 | 6 | 2.8451 | 6.29 | 0.0279 | Significant (p ≤ 0.05) |
| Col-II postwash in pH7 | Col-II 4 to 7 | 1.05074 | 1.01763 | 0.15298 | 0.1747 | 10 | 6 | 0.1426 | 11.76 | 0.889 | Not Significant (p > 0.0 |
| Col-I postwash in pH7 | Col-I 4 to 7 | 1.57403 | 1.14287 | 0.2551 | 0.0275 | 9 | 6 | 1.6804 | 8.19 | 0.1305 | Not Significant (p > 0.0 |

**Table S6.** Two-way ANOVA test results for shear-dependent compliance of collagen films are presented in Figure 3 (C) of the main text.

*Overall ANOVA*

| | DF | Sum of Squares | Mean Square | F Value | P Value |
|---|---|---|---|---|---|
| CollagenType | 1 | 1.05316 | 1.05316 | 3.67648 | 0.06544 |
| FormationpH | 1 | 0.38234 | 0.38234 | 1.33469 | 0.25774 |
| Model | 2 | 1.40387 | 0.70194 | 2.45038 | 0.10455 |
| Error | 28 | 8.02088 | 0.28646 | -- | -- |
| Corrected Total | 30 | 9.42475 | -- | -- | -- |

At the 0.05 level, the population means of **CollagenType** are **not significantly** different.
At the 0.05 level, the population means of **FormationpH** are **not significantly** different.

**Table S7**. Comparison of mechanical properties measured by AFM and QCM. The calculation of $E_{QCM}$ comes from assuming fully isotropic, homogeneous and linearly elastic films.

| | $E_{avg}$ [MPa] (AFM) | $J'_f$ [1/MPa] (QCM-D) | $G_f$ [MPa] ($1/J'_f$) | $E_{QCM}$ [MPa] (3x$G_f$) |
|---|---|---|---|---|
| Col-II pH 7 | 3.20 | 1.05 | 0.95 | 2.85 |
| Col-II pH 4 to 7 | 3.00 | 0.82 | 1.21 | 3.63 |
| Col-I pH 7 | 3.26 | 1.57 | 0.64 | 1.92 |
| Col-I pH 4 to 7 | 1.39 | 1.14 | 0.88 | 2.64 |

**Table S8**. Summary of QCM-D determined dSF and rLub film Sauerbrey masses before (pre) and after (post) PBS washes on collagen films.

| SAM - pH | dSF pre-wash $m_f$ [ng/cm$^2$] | dSF post-wash $m_f$ [ng/cm$^2$] | rLub pre-wash $m_f$ [ng/cm$^2$] | rLub post-wash $m_f$ [ng/cm$^2$] |
|---|---|---|---|---|
| Col-II - pH 7 | 300 $\pm$ 50 | 230 $\pm$ 50 | 50 $\pm$ 17 | 53 $\pm$ 18 |
| Col-II - pH 4 | 410 $\pm$ 60 | 340 $\pm$ 40 | 46 $\pm$ 7 | 50 $\pm$ 8 |
| Col-I – pH 7 | Little or no adsorption | Little or no adsorption | 31 $\pm$ 16 | 35 $\pm$ 18 |
| Col-I – pH 4 | 82 $\pm$ 4 | 17 $\pm$ 4 | 37 $\pm$ 14 | 41 $\pm$ 16 |

**Table S9.** *t*-test results for Sauerbrey mass of rLub and dSF films, presented in Figure 6 and the discussion of the main text.

| Group 1 | Group 2 | Mean 1 | Mean 2 | SEM 1 | SEM 2 | n1 | n2 | t-statistic | df | p-value | Result |
|---|---|---|---|---|---|---|---|---|---|---|---|
| R-LUB comparisons | | | | | | | | | | | |
| Lub Pre on Col I pH 7 | Lub Post on Col I pH 7 | 30.93333 | 34.66667 | 16.44611 | 17.96017 | 3 | 3 | -0.1533 | 3.97 | 0.8856 | Not Significant ($p > 0.05$) |
| Lub Pre on Col I pH 4 | Lub Post on Col I pH 4 | 37.16667 | 40.53333 | 14.1836 | 15.98649 | 3 | 3 | -0.1575 | 3.94 | 0.8826 | Not Significant ($p > 0.05$) |
| Lub Pre on Col II pH 7 | Lub Post on Col II pH 7 | 50.33333 | 52.7 | 17.13372 | 17.77254 | 3 | 3 | -0.0959 | 3.99 | 0.9282 | Not Significant ($p > 0.05$) |
| Lub Pre on Col II pH 4 | Lub Post on Col II pH 4 | 45.5 | 50.325 | 6.79853 | 7.50482 | 4 | 4 | -0.4765 | 5.94 | 0.6507 | Not Significant ($p > 0.05$) |
| Lub Pre on Col I pH 7 | Lub Pre on Col I pH 4 | 30.93333 | 37.16667 | 16.44611 | 14.1836 | 3 | 3 | -0.287 | 3.92 | 0.7886 | Not Significant ($p > 0.05$) |
| Lub Post on Col I pH 7 | Lub Post on Col I pH 4 | 34.66667 | 40.53333 | 17.96017 | 15.98649 | 3 | 3 | -0.244 | 3.95 | 0.8194 | Not Significant ($p > 0.05$) |
| Lub Pre on Col II pH 7 | Lub Pre on Col II pH 4 | 50.33333 | 45.5 | 17.13372 | 6.79853 | 3 | 4 | 0.2622 | 2.64 | 0.8123 | Not Significant ($p > 0.05$) |
| Lub Post on Col II pH 7 | Lub Post on Col II pH 4 | 52.7 | 50.325 | 17.77254 | 7.50482 | 3 | 4 | 0.1231 | 2.72 | 0.9105 | Not Significant ($p > 0.05$) |
| Lub Pre on Col I pH 7 | Lub Pre on Col II pH 7 | 30.93333 | 50.33333 | 16.44611 | 17.13372 | 3 | 3 | -0.8169 | 3.99 | 0.4599 | Not Significant ($p > 0.05$) |
| Lub Post on Col I pH 7 | Lub Post on Col II pH 7 | 34.66667 | 52.7 | 17.96017 | 17.77254 | 3 | 3 | -0.7137 | 4 | 0.5148 | Not Significant ($p > 0.05$) |
| Lub Pre on Col I pH 4 | Lub Pre on Col II pH 4 | 37.16667 | 45.5 | 14.1836 | 6.79853 | 3 | 4 | -0.5298 | 2.92 | 0.6339 | Not Significant ($p > 0.05$) |
| Lub Post on Col I pH 4 | Lub Post on Col II pH 4 | 40.53333 | 50.325 | 15.98649 | 7.50482 | 3 | 4 | -0.5544 | 2.89 | 0.6193 | Not Significant ($p > 0.05$) |
| dSF comparisons | | | | | | | | | | | |
| dSF prewash on Col-II in pH 7 | dSF postwash on Col-II in pH 7 | 299.537 | 227.1167 | 48.13111 | 27.11011 | 3 | 3 | 1.311 | 3.15 | 0.2772 | Not Significant ($p > 0.05$) |
| dSF prewash on Col-I in pH 7 | dSF postwash on Col-I in pH 7 | 5.54752 | -57.7956 | 18.04803 | 37.63371 | 4 | 4 | 1.5177 | 4.31 | 0.1987 | Not Significant ($p > 0.05$) |
| dSF prewash on Col-II in pH 4 | dSF postwash on Col-II in pH 4 | 414.3653 | 340.1049 | 62.09944 | 43.96293 | 3 | 3 | 0.976 | 3.6 | 0.39 | Not Significant ($p > 0.05$) |
| dSF prewash on Col-I in pH 4 | dSF postwash on Col-I in pH 4 | 82.36258 | 16.47167 | 4.21024 | 3.62467 | 3 | 3 | 11.8603 | 3.91 | 0.0003 | Significant ($p \le 0.05$) |
| dSF prewash on Col-II in pH 7 | dSF prewash on Col-I in pH 7 | 299.537 | 5.54752 | 48.13111 | 18.04803 | 3 | 4 | 5.7192 | 2.57 | 0.016 | Significant ($p \le 0.05$) |
| dSF postwash on Col-II in pH 7 | dSF postwash on Col-I in pH 7 | 227.1167 | -57.7956 | 27.11011 | 37.63371 | 3 | 4 | 6.1428 | 4.93 | 0.0017 | Significant ($p \le 0.05$) |
| dSF prewash on Col-II in pH 4 | dSF prewash on Col-I in pH 4 | 414.3653 | 82.36258 | 62.09944 | 4.21024 | 3 | 3 | 5.3341 | 2.02 | 0.0327 | Significant ($p \le 0.05$) |
| dSF postwash on Col-II in pH 4 | dSF postwash on Col-I in pH 4 | 340.1049 | 16.47167 | 43.96293 | 3.62467 | 3 | 3 | 7.3366 | 2.03 | 0.0174 | Significant ($p \le 0.05$) |
| dSF prewash on Col-II in pH 7 | dSF prewash on Col-II in pH 4 | 299.537 | 414.3653 | 48.13111 | 62.09944 | 3 | 3 | -1.4615 | 3.77 | 0.2219 | Not Significant ($p > 0.05$) |
| dSF postwash on Col-II in pH 7 | dSF postwash on Col-II in pH 4 | 227.1167 | 340.1049 | 27.11011 | 43.96293 | 3 | 3 | -2.1876 | 3.33 | 0.1076 | Not Significant ($p > 0.05$) |
| dSF prewash on Col-I in pH 7 | dSF prewash on Col-I in pH 4 | 5.54752 | 82.36258 | 18.04803 | 4.21024 | 4 | 3 | -4.1449 | 3.32 | 0.0209 | Significant ($p \le 0.05$) |
| dSF postwash on Col-I in pH 7 | dSF postwash on Col-I in pH 4 | -57.7956 | 16.47167 | 37.63371 | 3.62467 | 4 | 3 | -1.9643 | 3.06 | 0.1426 | Not Significant ($p > 0.05$) |
| dSF vs R-LUB comparisons | | | | | | | | | | | |
| dSF prewash on Col-II in pH 7 | Lub Pre on Col II pH 7 | 299.537 | 50.33333 | 48.13111 | 17.13372 | 3 | 3 | 4.8778 | 2.5 | 0.0249 | Significant ($p \le 0.05$) |
| dSF postwash on Col-II in pH 7 | Lub Post on Col II pH 7 | 227.1167 | 52.7 | 27.11011 | 17.77254 | 3 | 3 | 5.3805 | 3.45 | 0.0087 | Significant ($p \le 0.05$) |
| dSF prewash on Col-I in pH 7 | Lub Pre on Col I pH 7 | 5.54752 | 30.93333 | 18.04803 | 16.44611 | 4 | 3 | -1.0397 | 4.94 | 0.3467 | Not Significant ($p > 0.05$) |
| dSF postwash on Col-I in pH 7 | Lub Post on Col I pH 7 | -57.7956 | 34.66667 | 37.63371 | 17.96017 | 4 | 3 | -2.2173 | 4.2 | 0.0877 | Not Significant ($p > 0.05$) |
| dSF prewash on Col-II in pH 4 | Lub Pre on Col II pH 4 | 414.3653 | 45.5 | 62.09944 | 6.79853 | 3 | 4 | 5.9046 | 2.05 | 0.026 | Significant ($p \le 0.05$) |
| dSF postwash on Col-II in pH 4 | Lub Post on Col II pH 4 | 340.1049 | 50.325 | 43.96293 | 7.50482 | 3 | 4 | 6.4975 | 2.12 | 0.0198 | Significant ($p \le 0.05$) |
| dSF prewash on Col-I in pH 4 | Lub Pre on Col I pH 4 | 82.36258 | 37.16667 | 4.21024 | 14.1836 | 3 | 3 | 3.0547 | 2.35 | 0.0756 | Not Significant ($p > 0.05$) |
| dSF postwash on Col-I in pH 4 | Lub Post on Col I pH 4 | 16.47167 | 40.53333 | 3.62467 | 15.98649 | 3 | 3 | -1.4679 | 2.21 | 0.2687 | Not Significant ($p > 0.05$) |

**Table S10.** Two-way ANOVA test results for Sauerbrey mass of dSF on collagen films, presented in Figure 6 (A).

*Overall ANOVA*

| | DF | Sum of Squares | Mean Square | F Value | P Value |
|---|---|---|---|---|---|
| CollagenType | 1 | 295065.40195 | 295065.40195 | 86.07504 | 3.14624E-6 |
| Formation pH | 1 | 27405.39375 | 27405.39375 | 7.99457 | 0.01793 |
| Model | 2 | 337036.7023 | 168518.35115 | 49.15935 | 6.7063E-6 |
| Error | 10 | 34280.01872 | 3428.00187 | -- | -- |
| Corrected Total | 12 | 371316.72102 | -- | -- | -- |

At the 0.05 level, the population means of **CollagenType** are **significantly** different.
At the 0.05 level, the population means of **Formation pH** are **significantly** different.

**Table S11.** Two-way ANOVA test results for Sauerbrey mass of rLub on collagen films, presented in Figure 6 (B).

*Overall ANOVA*

| | DF | Sum of Squares | Mean Square | F Value | P Value |
|---|---|---|---|---|---|
| CollagenType | 1 | 597.82173 | 597.82173 | 0.98097 | 0.34532 |
| Formation pH | 1 | 6.95625 | 6.95625 | 0.01141 | 0.91703 |
| Model | 2 | 617.13911 | 308.56956 | 0.50633 | 0.61736 |
| Error | 10 | 6094.20089 | 609.42009 | -- | -- |
| Corrected Total | 12 | 6711.34 | -- | -- | -- |

At the 0.05 level, the population means of **CollagenType** are **not significantly** different.
At the 0.05 level, the population means of **Formation pH** are **not significantly** different.

**Table S12**. Summary of shear-dependent compliance determined for dSF and rLub and their respective collagen films (as represented in Figure 7 (B)) before (pre) and after (post) PBS washes.

| Collagen-pH | dSF pre-wash $J'_{\mathrm{f}}$ [1/MPa] | dSF post-wash $J'_{\mathrm{f}}$ [1/MPa] | rLub pre-wash $J'_{\mathrm{f}}$ [1/MPa] | rLub post-wash $J'_{\mathrm{f}}$ [1/MPa] |
|---|---|---|---|---|
| Col-II - pH 7 | 1.18 ± 0.0 | 1.14 ± 0.34 | 0.96 ± 0.12 | 0.96 ± 0.12 |
| Col-II - pH 4 | 0.78 ± 0.22 | 0.72 ± 0.28 | 1.08 ± 0.14 | 1.08 ± 0.15 |
| Col-I – pH 7 | 2.11± 0.20 | 1.97 ± 0.23 | 1.02 ± 0.05 | 1.02 ± 0.05 |
| Col-I – pH 4 | 1.38 ± 0.07 | 1.40 ± 0.06 | 1.11 ± 0.03 | 1.12 ± 0.04 |

**Table S13.** *t*-test results for shear dependent compliance of the stacked rLub and dSF films with collagen, presented in Figure 7 of the main text.

dSF vs Collagen

| Group 1 | Group 2 | Mean 1 | Mean 2 | SEM 1 | SEM 2 | n1 | n2 | t-statistic | df | p-value | Result |
|---|---|---|---|---|---|---|---|---|---|---|---|
| dSF+BSA_Col-II pH 7 prewash | dSF+BSA_Col-II pH 7 postwash | 1.1758 | 1.13758 | 0.30102 | 0.33852 | 3 | 3 | 0.0844 | 3.95 | 0.9369 | Not Significant (p > 0.05) |
| dSF+BSA_ Col-II pH 4 postwash | dSF+BSA_ Col-II pH 4 prewash | 0.72166 | 0.77516 | 0.2844 | 0.22289 | 3 | 3 | -0.1481 | 3.78 | 0.8898 | Not Significant (p > 0.05) |
| dSF+BSA on Col-I in pH4 prewash | dSF+BSA on Col-I in pH4 postwash | 1.38323 | 1.39505 | 0.0701 | 0.0645 | 3 | 3 | -0.1241 | 3.97 | 0.9073 | Not Significant (p > 0.05) |
| dSF+BSA_Col-II pH 7 prewash | dSF+BSA_ Col-II pH 4 prewash | 1.1758 | 0.77516 | 0.30102 | 0.22289 | 3 | 3 | 1.0696 | 3.69 | 0.3498 | Not Significant (p > 0.05) |
| dSF+BSA_Col-II pH 7 postwash | dSF+BSA_ Col-II pH 4 postwash | 1.13758 | 0.72166 | 0.33852 | 0.2844 | 3 | 3 | 0.9407 | 3.88 | 0.4016 | Not Significant (p > 0.05) |
| dSF+BSA_ Col-II pH 4 postwash | dSF+BSA on Col-I in pH4 postwash | 0.72166 | 1.39505 | 0.2844 | 0.0645 | 3 | 3 | -2.3091 | 2.21 | 0.1354 | Not Significant (p > 0.05) |
| dSF+BSA_ Col-II pH 4 prewash | dSF+BSA on Col-I in pH4 prewash | 0.77516 | 1.38323 | 0.22289 | 0.0701 | 3 | 3 | -2.6024 | 2.39 | 0.1012 | Not Significant (p > 0.05) |
| Col-II 4 to 7 | dSF+BSA_ Col-II pH 4 prewash | 1.01763 | 0.77516 | 0.1747 | 0.22289 | 6 | 3 | 0.8562 | 4.53 | 0.4348 | Not Significant (p > 0.05) |
| Col-II 4 to 7 | dSF+BSA_ Col-II pH 4 postwash | 1.01763 | 0.72166 | 0.1747 | 0.2844 | 6 | 3 | 0.8867 | 3.59 | 0.4306 | Not Significant (p > 0.05) |
| Col-I 4 to 7 | dSF+BSA on Col-I in pH4 prewash | 1.14287 | 1.38323 | 0.0275 | 0.0701 | 6 | 3 | -3.192 | 2.64 | 0.0593 | Not Significant (p > 0.05) |
| Col-I 4 to 7 | dSF+BSA on Col-I in pH4 postwash | 1.14287 | 1.39505 | 0.0275 | 0.0645 | 6 | 3 | -3.5965 | 2.76 | 0.0422 | Significant (p ≤ 0.05) |
| Col-II postwash in pH7 | dSF+BSA_Col-II pH 7 prewash | 1.05074 | 1.1758 | 0.15298 | 0.30102 | 10 | 3 | -0.3704 | 3.12 | 0.7348 | Not Significant (p > 0.05) |
| Col-II postwash in pH7 | dSF+BSA_Col-II pH 7 postwash | 1.05074 | 1.13758 | 0.15298 | 0.33852 | 10 | 3 | -0.2338 | 2.87 | 0.8308 | Not Significant (p > 0.05) |
| dSF+BSA on Col-I in pH7 prewash | Col-I postwash in pH7 | 2.11049 | 1.57403 | 0.19878 | 0.2551 | 4 | 9 | 1.6588 | 10.42 | 0.1269 | Not Significant (p > 0.05) |
| dSF+BSA on Col-I in pH7 postwash | Col-I postwash in pH7 | 1.96525 | 1.57403 | 0.23355 | 0.2551 | 4 | 9 | 1.1311 | 9.41 | 0.286 | Not Significant (p > 0.05) |
| dSF+BSA on Col-I in pH7 prewash | dSF+BSA on Col-I in pH7 postwash | 2.11049 | 1.96525 | 0.19878 | 0.23355 | 4 | 4 | 0.4736 | 5.85 | 0.653 | Not Significant (p > 0.05) |

LUB vs Collagen

| Group 1 | Group 2 | Mean 1 | Mean 2 | SEM 1 | SEM 2 | n1 | n2 | t-statistic | df | p-value | Result |
|---|---|---|---|---|---|---|---|---|---|---|---|
| Jf Lub Pre on Col I pH 7 | Col-I postwash in pH7 | 1.01962 | 1.57403 | 0.04665 | 0.2551 | 3 | 9 | -2.1379 | 8.51 | 0.063 | Not Significant (p > 0.05) |
| Jf Lub Post on Col I pH 7 | Col-I postwash in pH7 | 1.01646 | 1.57403 | 0.0461 | 0.2551 | 3 | 9 | -2.1509 | 8.49 | 0.0617 | Not Significant (p > 0.05) |
| Jf Lub Pre on Col I pH 4 | Col-I 4 to 7 | 1.11492 | 1.14287 | 0.03475 | 0.0275 | 3 | 6 | -0.6307 | 4.57 | 0.5584 | Not Significant (p > 0.05) |
| JfLub Post on Col I pH 4 | Col-I 4 to 7 | 1.11547 | 1.14287 | 0.03618 | 0.0275 | 3 | 6 | -0.6029 | 4.39 | 0.5763 | Not Significant (p > 0.05) |
| Jf Lub Pre on Col II pH 7 | Col-II postwash in pH7 | 0.95644 | 1.05074 | 0.11953 | 0.15298 | 3 | 10 | -0.4857 | 8.72 | 0.6391 | Not Significant (p > 0.05) |
| Jf Lub Post on Col II pH 7 | Col-II postwash in pH7 | 0.95979 | 1.05074 | 0.11768 | 0.15298 | 3 | 10 | -0.4712 | 8.85 | 0.6489 | Not Significant (p > 0.05) |
| Jf Lub Pre on Col II pH 4 | Col-II 4 to 7 | 1.0778 | 1.01763 | 0.14402 | 0.1747 | 4 | 6 | 0.2658 | 7.97 | 0.7972 | Not Significant (p > 0.05) |
| Jf Lub Post on Col II pH 4 | Col-II 4 to 7 | 1.08184 | 1.01763 | 0.1463 | 0.1747 | 4 | 6 | 0.2818 | 7.95 | 0.7853 | Not Significant (p > 0.05) |
| Jf Lub Pre on Col I pH 7 | Jf Lub Pre on Col I pH 4 | 1.01962 | 1.11492 | 0.04665 | 0.03475 | 3 | 3 | -1.6383 | 3.7 | 0.1825 | Not Significant (p > 0.05) |
| Jf Lub Post on Col I pH 7 | JfLub Post on Col I pH 4 | 1.01646 | 1.11547 | 0.0461 | 0.03618 | 3 | 3 | -1.6895 | 3.79 | 0.1704 | Not Significant (p > 0.05) |
| Jf Lub Pre on Col II pH 7 | Jf Lub Pre on Col II pH 4 | 0.95644 | 1.0778 | 0.11953 | 0.14402 | 3 | 4 | -0.6484 | 5 | 0.5453 | Not Significant (p > 0.05) |
| Jf Lub Post on Col II pH 7 | Jf Lub Post on Col II pH 4 | 0.95979 | 1.08184 | 0.11768 | 0.1463 | 3 | 4 | -0.65 | 5 | 0.5443 | Not Significant (p > 0.05) |
| Jf Lub Pre on Col I pH 7 | Jf Lub Pre on Col II pH 7 | 1.01962 | 0.95644 | 0.04665 | 0.11953 | 3 | 3 | 0.4924 | 2.6 | 0.661 | Not Significant (p > 0.05) |
| Jf Lub Post on Col I pH 7 | Jf Lub Post on Col II pH 7 | 1.01646 | 0.95979 | 0.0461 | 0.11768 | 3 | 3 | 0.4484 | 2.6 | 0.6886 | Not Significant (p > 0.05) |
| Jf Lub Pre on Col I pH 4 | Jf Lub Pre on Col II pH 4 | 1.11492 | 1.0778 | 0.03475 | 0.14402 | 3 | 4 | 0.2506 | 3.34 | 0.8168 | Not Significant (p > 0.05) |
| Jf Lub Post on Col I pH 4 | Jf Lub Post on Col II pH 4 | 1.11547 | 1.08184 | 0.03618 | 0.1463 | 3 | 4 | 0.2231 | 3.36 | 0.8363 | Not Significant (p > 0.05) |

dSF vs LUB on collagen

| Group 1 | Group 2 | Mean 1 | Mean 2 | SEM 1 | SEM 2 | n1 | n2 | t-statistic | df | p-value | Result |
|---|---|---|---|---|---|---|---|---|---|---|---|
| Jf Lub Post on Col II pH 4 | dSF+BSA_ Col-II pH 4 postwash | 1.08184 | 0.72166 | 0.1463 | 0.2844 | 4 | 3 | 1.1262 | 3.06 | 0.3407 | Not Significant (p > 0.05) |
| Jf Lub Pre on Col I pH 7 | dSF+BSA on Col-I in pH7 prewash | 1.01962 | 1.96525 | 0.04665 | 0.23355 | 3 | 4 | -3.9705 | 3.24 | 0.0248 | Significant (p ≤ 0.05) |
| Jf Lub Post on Col I pH 7 | dSF+BSA on Col-I in pH7 postwash | 1.01646 | 2.11049 | 0.0461 | 0.19878 | 3 | 4 | -5.3614 | 3.32 | 0.0098 | Significant (p ≤ 0.05) |
| Jf Lub Pre on Col I pH 4 | dSF+BSA on Col-I in pH4 prewash | 1.11492 | 1.38323 | 0.03475 | 0.0701 | 3 | 3 | -3.4293 | 2.93 | 0.0432 | Significant (p ≤ 0.05) |
| JfLub Post on Col I pH 4 | dSF+BSA on Col-I in pH4 postwash | 1.11547 | 1.39505 | 0.03618 | 0.0645 | 3 | 3 | -3.7804 | 3.15 | 0.0299 | Significant (p ≤ 0.05) |
| Jf Lub Pre on Col II pH 7 | dSF+BSA_Col-II pH 7 prewash | 0.95644 | 1.1758 | 0.11953 | 0.30102 | 3 | 3 | -0.6773 | 2.62 | 0.5532 | Not Significant (p > 0.05) |
| Jf Lub Post on Col II pH 7 | dSF+BSA_Col-II pH 7 postwash | 0.95979 | 1.13758 | 0.11768 | 0.33852 | 3 | 3 | -0.4961 | 2.48 | 0.6605 | Not Significant (p > 0.05) |
| Jf Lub Pre on Col II pH 4 | dSF+BSA_ Col-II pH 4 prewash | 1.0778 | 0.77516 | 0.14402 | 0.22289 | 4 | 3 | 1.1404 | 3.6 | 0.3242 | Not Significant (p > 0.05) |
| Jf Lub Post on Col II pH 4 | dSF+BSA_ Col-II pH 4 postwash | 1.08184 | 0.72166 | 0.1463 | 0.2844 | 4 | 3 | 1.1262 | 3.06 | 0.3407 | Not Significant (p > 0.05) |

### REFERENCES


[1] S. T. Ahmed, D. R. Jaramillo Pinto, L. Vitkova, U. Honey, W. Flores, K. L. Lunny, K. A. Cutter, Y. Wen, K. De France, R. C. Andresen Eguiluz, *Colloids Surf. B Biointerfaces* **2026**, *258*, 115224.
